# Yeast adapts to diverse ecological niches driven by genomics and metabolic reprogramming

**DOI:** 10.1101/2024.06.14.598782

**Authors:** Haoyu Wang, Jens Nielsen, Yongjin Zhou, Hongzhong Lu

**Author notes:** Correspondence (H.Z. Lu), (Y.Z. Zhou).

## Abstract

The famous model organism - *Saccharomyces cerevisiae* is widely present in a variety of natural and human-associated habitats. Despite extensive studies of this organism, the metabolic mechanisms driving its adaptation to varying niches remain elusive. We here gathered genomic resources from 1,807 *S. cerevisiae* strains and assembled them into a high-quality pan-genome, facilitating the comprehensive characterization of genetic diversity across isolates. Utilizing the pan-genome, 1,807 strain-specific genome-scale metabolic models (ssGEMs) were generated, which performed well in quantitative predictions of cellular phenotypes, thus helping to examine the metabolic disparities among all *S. cerevisiae* strains. Integrative analyses of fluxomic and transcriptomics with ssGEMs showcased the ubiquitous transcriptional regulation in certain metabolic sub-pathways (i.e., amino acid synthesis) at a population level. Additionally, the gene/reaction loss analysis through the ssGEMs refined by transcriptomics showed that *S. cerevisiae* strains from various ecological niches had undergone reductive evolution at both the genomic and metabolic network levels when compared to wild isolates. Finally, the compiled analyses of the pan-genome, transcriptome, and metabolic fluxome revealed remarkable metabolic differences among *S. cerevisiae* strains originating from distinct oxygen-limited niches, including human gut and cheese environments, and identified convergent metabolic evolution, such as downregulation of oxidative phosphorylation pathways. Together, these results illustrate how yeast adapts to distinct niches modulated by genomic and metabolic reprogramming, and provide computational resources for translating yeast genotype to fitness in future studies.

## Introduction

As one of the most famous eukaryotic model organisms, the budding yeast *Saccharomyces cerevisiae*, has been widely used in areas of synthetic and systems biology^1–5^. The first whole genome sequence of *S. cerevisiae* was released in 1996, followed by numerous molecular studies using its laboratory mutants^3,6–8^. Well adapted to laboratory environments, *S. cerevisiae* is also ubiquitous in both natural and human-associated niches. The wide distribution around the earth has shaped the genetic and phenotypic diversity of distinct *S. cerevisiae* isolates^9^. Elucidating how *S. cerevisiae* strains adapt to varied environments could provide novel insights into the metabolic rewiring of eukaryotic organisms driven by natural and artificial selection^10^.

The advancement of high-throughput sequencing technology has made it possible to profile the genomic evolution of *S. cerevisiae* on a larger scale. Over the past decade, amounts of genome sequencing have been performed to characterize the genetic diversity of *S. cerevisiae* strains sampled from a wide range of habitats, such as wine, plants and human-guts^11–17^. Notably, Peter et al. reported the whole genome sequencing for 1,011 *S. cerevisiae* strains worldwide, successfully delineating the global evolutionary portrait of the species^9^. More recently, the same group performed pan-transcriptome analysis of ∼1,000 *S. cerevisiae* strains and illustrated the gene expression patterns across the major evolutionary clades^18^. Together, these omics datasets provide valuable resources to study the evolution of *S. cerevisiae* at system and population levels.

The pan-genome, defined as the collection of all genes encompassed by a group of individuals from a certain species, has become as a powerful tool to characterize the genomic and metabolic diversity of studied species^19^. Currently, there exist two representative pan-genomes for *S. cerevisiae*, here named as Sce-pan1011 and Sce-pan1364, which were built for 1,011 and 1,364 isolates, respectively. By contrast, the Sce-pan1011 contains a total of 7,796 ORFs, while the Sce-pan1364 has only 7,078 ORFs^9,20^. With the accumulation of newly sequenced *S. cerevisiae* strains, it is thus critical to re-build the pan-genome of *S. cerevisiae* considering all sequenced isolates, as well as refining its architecture to solve the existing conflict between the aforementioned two versions of pan-genomes.

Genome-scale metabolic models (GEMs) provide a mathematical framework representing the entire metabolism of an organism^21^. It could be served as a tool to mechanistically link genotype to phenotype. Since the first GEM published in 2003, *S. cerevisiae* GEMs have been continuously improved through iterative updates by the community^22,23^. During GEMs reconstruction, the genomic information of *S. cerevisiae* S288c was intensively leveraged, while omitting the gene gain and loss existing in other isolates. Therefore, it is challenging to directly apply the current *S. cerevisiae* GEMs-Yeast8 to simulate the metabolic diversity of other isolates. In this regard, Lu et al have built pan-GEMs for 1,011 *S. cerevisiae* and, for the first time, developed strain-specific GEMs for all the studied strains, which illustrating metabolic conservation and variation at single-strain resolution. However, the limitation of omics datasets, i.e., transcriptomics and phenomics, to some extent, hinders the applications of GEMs in reflecting the in vivo metabolic diversity of those strains.

To systematically elucidate the molecular mechanisms underpinning the adaptation of *S. cerevisiae* strains across a variety of environments, herein we reconstructed a new version of pan-genome for *S. cerevisiae* utilizing the high-quality genome sequences from 1,807 isolates, which helps to comprehensively describe genetic diversity within the species. Subsequently, inspired by the detailed function annotation of the newly built pan-genome, 1,807 strain-specific genome-scale metabolic models (ssGEMs) were established to quantitatively assess metabolic differences between strains. We further integrated large-scale transcriptomics data with ssGEMs to investigate the transcriptional regulation of metabolic network at the population level. Finally, a multidimensional integration analysis of the pan-genome, transcriptome, and fluxome was performed to characterize the genetic and metabolic features of *S. cerevisiae* strains that originated from anaerobic environments, including the human gut and dairy niches. Overall, with advanced genomic and modelling analysis, our work shows how the metabolism of *S. cerevisiae* was dynamically reprogrammed to adapt to diverse ecological niches around the world.

## Results

### Pan-genome of 1,807 *S. cerevisiae* strains

To comprehensively investigate the genetic diversity of *S. cerevisiae*, we collected 1,913 assembled genome sequences of *S. cerevisiae* strains from the National Center for Biotechnology Information (NCBI) and published literatures^9,11,13,14,16,17,20^. After performing unified genome annotation and stringent quality control, we selected 1,807 high-quality annotated genomes to construct the pan-genome (Supplementary Fig. 1, Material and Methods). We also refined the sampled environments for each strain used in the following analysis. Our collection, curation, and refinement of each isolate’s genome laid a solid foundation for the reconstruction of the *S. cerevisiae* pan-genome from scratch. Ultimately, a total of 7,514 distinct ORFs were identified throughout the 1,807 genomes to construct the new version of the *S. cerevisiae* pan-genome, which was named as Sce-pan1807 (Fig. 1a, Supplementary Fig. 1). The size of our pan-genome could be comparable with two previously created pan-genomes - Sce- pan1011 and Sce-pan1364^9,20^, which possess 7,796 and 7,078 ORFs, respectively (Table 1). During *S. cerevisiae* pan-genome reconstruction, we carefully optimized the whole pipeline and tuned the criteria during the gene cluster analysis to guarantee the high quality of this new- version pan-genome.

**Figure 1.**
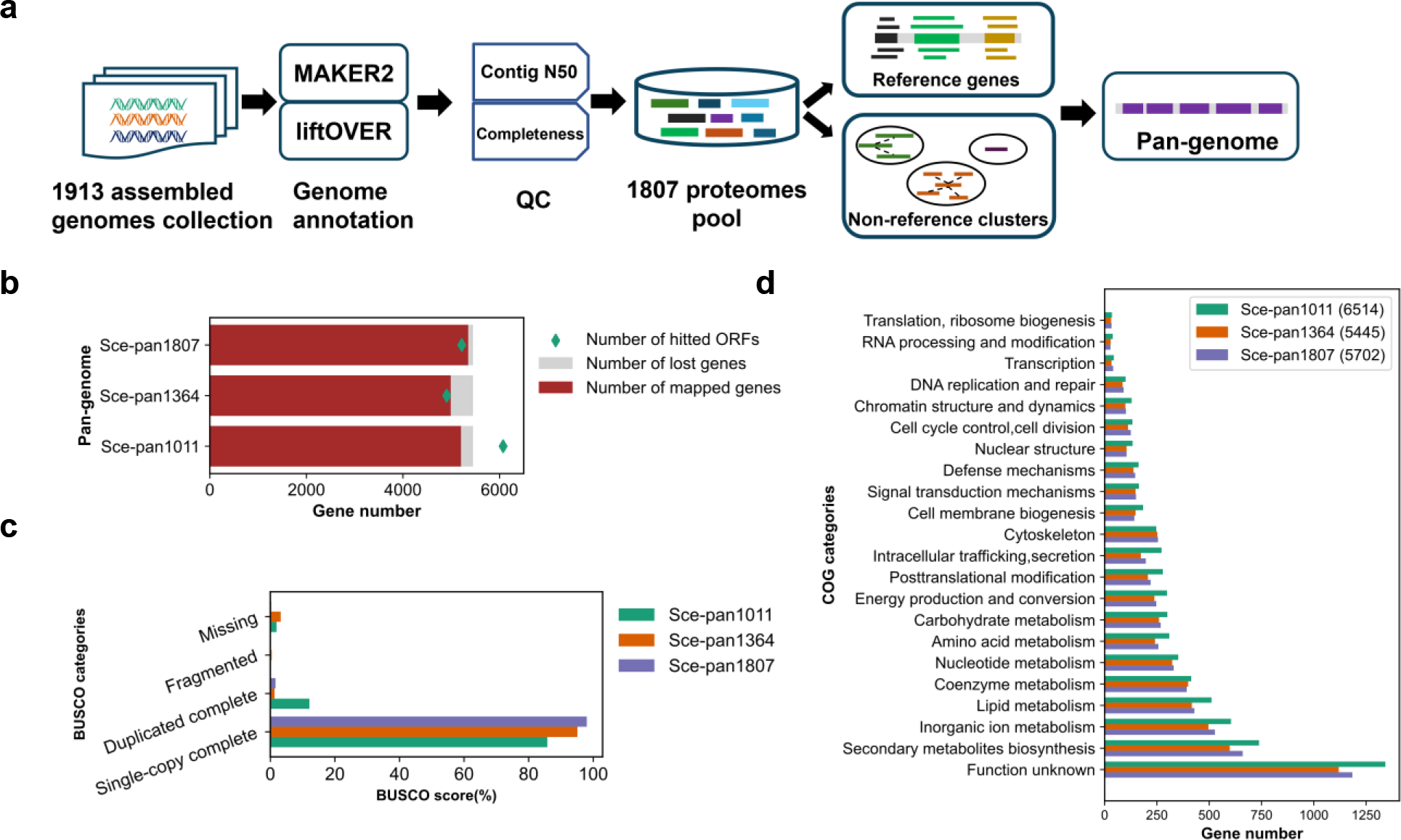
Construction and quality analysis of the *S. cerevisiae* pan-genome. Schematic overview of the workflow employed for constructing the *S. cerevisiae* pan-genome (a). Comparative analysis in gene representation between the newly built pan-genome (Sce- pan1807) and the previously two published ones (Sce-pan1011, Sce-pan1364). During the mapping analysis, the genome of *S. cerevisiae* CEN.PK was set as the reference genome. ‘Number of hitted ORFs’ indicates the number of pan genes that have been detected in the reference genome. ‘Number of lost genes’ indicates the number of reference strain’s genes that have not been detected in the pan-genome. ‘Number of mapped genes’ indicates the number of the reference strain’s genes that have been detected in pan-genome (b). Assessment of pan- genome completeness using the BUSCO method and the saccharomycetes_odb10 database (c). Functional categorization of pan-genomes using Clusters of Orthologous Groups (COG) (d). The number in brackets in the legend indicates the number of functionally annotated genes in the pan-genomes.

**Table 1.**
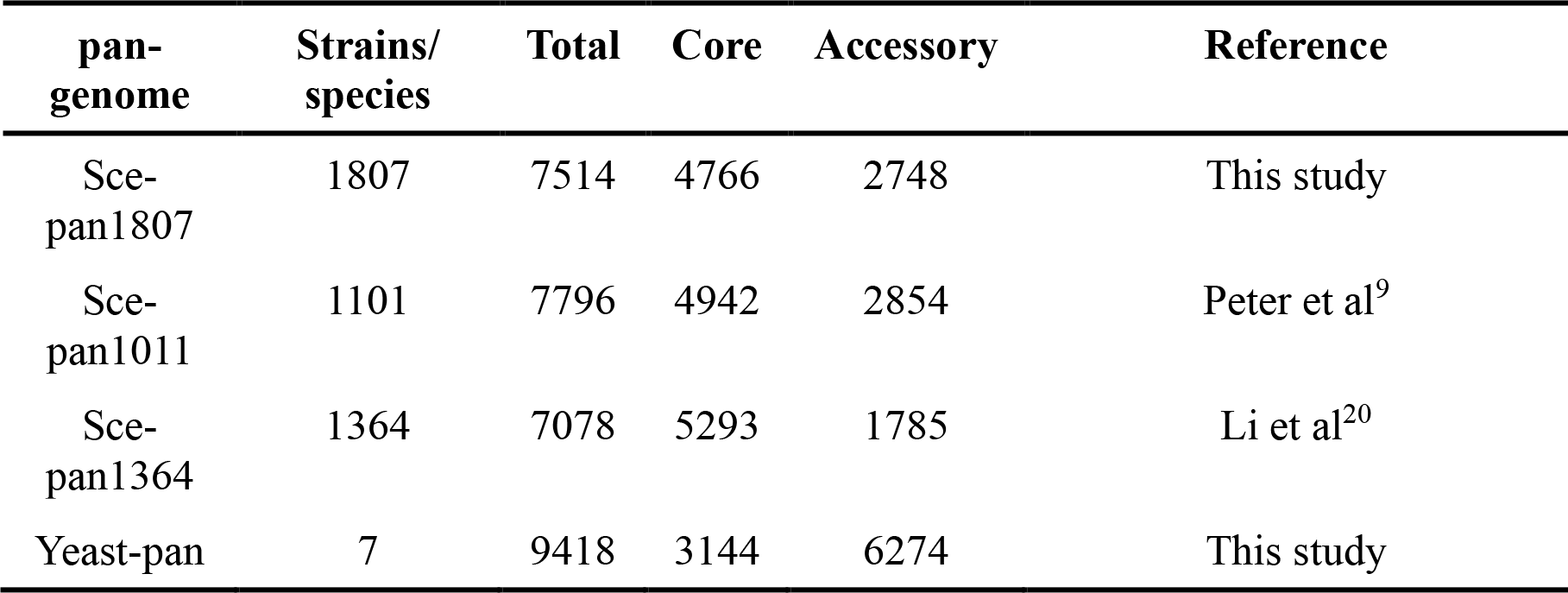
Comparison of all pan-genomes used in this study.

Given that the pan-genome was defined as the collective of all genes from a species, we first evaluated the capacity of our pan-genome and the previous ones in representing the genomic information from two well-studied strains. Genomes from *S. cerevisiae* CEN.PK and S288c were employed to map against each pan-genome’s representative sequences by sequence alignment (Fig. 1b, Supplementary Fig. 1d). All 3 pan-genomes could cover most genes in CEN.PK (∼98% for Sce-pan1807, ∼ 95% for Sce-pan1011 and ∼91% for Sce-pan1364); however, each pan-genome still exhibited some degree of gene missing during the above blast analysis. Particularly, the prior pan-genome Sce-pan1364, containing 7,078 genes, was accompanied with the highest number of missing genes. 249 genes from CEN.PK could not be mapped onto the Sce-pan1364, indicating that 7,078 ORFs in the Sce-pan1364 are insufficient to represent all genes in the species. By comparison, for Sce-pan1011, the hitted pan-gene number (6,075) is unreasonably larger than the total gene number of CEN.PK (5,451), implying that Sce-pan1011 may contain some redundant genes (Fig. 1b). The same analysis for S288c showed consistent results (Supplementary Fig. 1d). The quality of the three pan-genomes was further corroborated by the genome completeness evaluation using BUSCO^24^. All 3 pan- genomes exhibited nearly no fragmented genes. However, the pan-genome Sce-pan1364 has the highest missing score, while Sce-pan1011 was accompanied with the highest duplicated score (Fig. 1c). Collectively, these analyses indicate that our pan-genome performs well in reflecting the genomic information from different *S. cerevisiae* strains.

To illustrate the metabolic functions encoded in pan-genome, we performed the functional annotation for all 3 pan-genomes utilizing the eggNOG tool^25^, as well as the functional enrichment analysis based on the annotation from Clusters of Orthologous Groups (COGs). We discovered that the functional distribution of COG categories throughout the 3 pan-genomes displays a similar tendency (Fig. 1d), albeit there were some slight deviations across all COG categories, which may be due to the differences in the coverage and quality scores of the 3 pan- genomes.

### Accurate definition of core genome

The core genome is a collection of genes present in most strains of a given species, thus representing the shared genetic information from the studied species. To accurately define the core genome of *S. cerevisiae*, we performed a thorough analysis of the core genome size considering the frequency of ORFs occurring in different strains (Fig. 2a). We found that as the percentage threshold defining the core genes increasing from 95% to 100%, the core genome size decreased from 5,496 to 939. Notably, when the threshold surpassed 99%, the core genome size experienced a precipitous decline. The rapid decline after 99% could be primarily due to errors and incompleteness in the sequencing and assembly processes used to generate the large-scale genome dataset (1,807 genomes) collected here, which is a common issue in many population genomics studies^26–28^. To circumvent this, we selected 99% as the threshold to define the core genome in this study, yielding 4,766 core genes and 2,748 accessory genes within our *S. cerevisiae* pan-genome. By comparison, the core genome size is smaller in our work than that in the previous pan-genome assemblies (Table 1). The slightly larger core genome size in Sce-pan1011 (4,942) could be due to the redundancy discussed above (Fig. 1b, c), while the significantly larger size in Sce-pan1364 (5,293) might be due to the fact that a more relaxed threshold (95%) was utilized to define core genes^20^.

**Figure 2.**
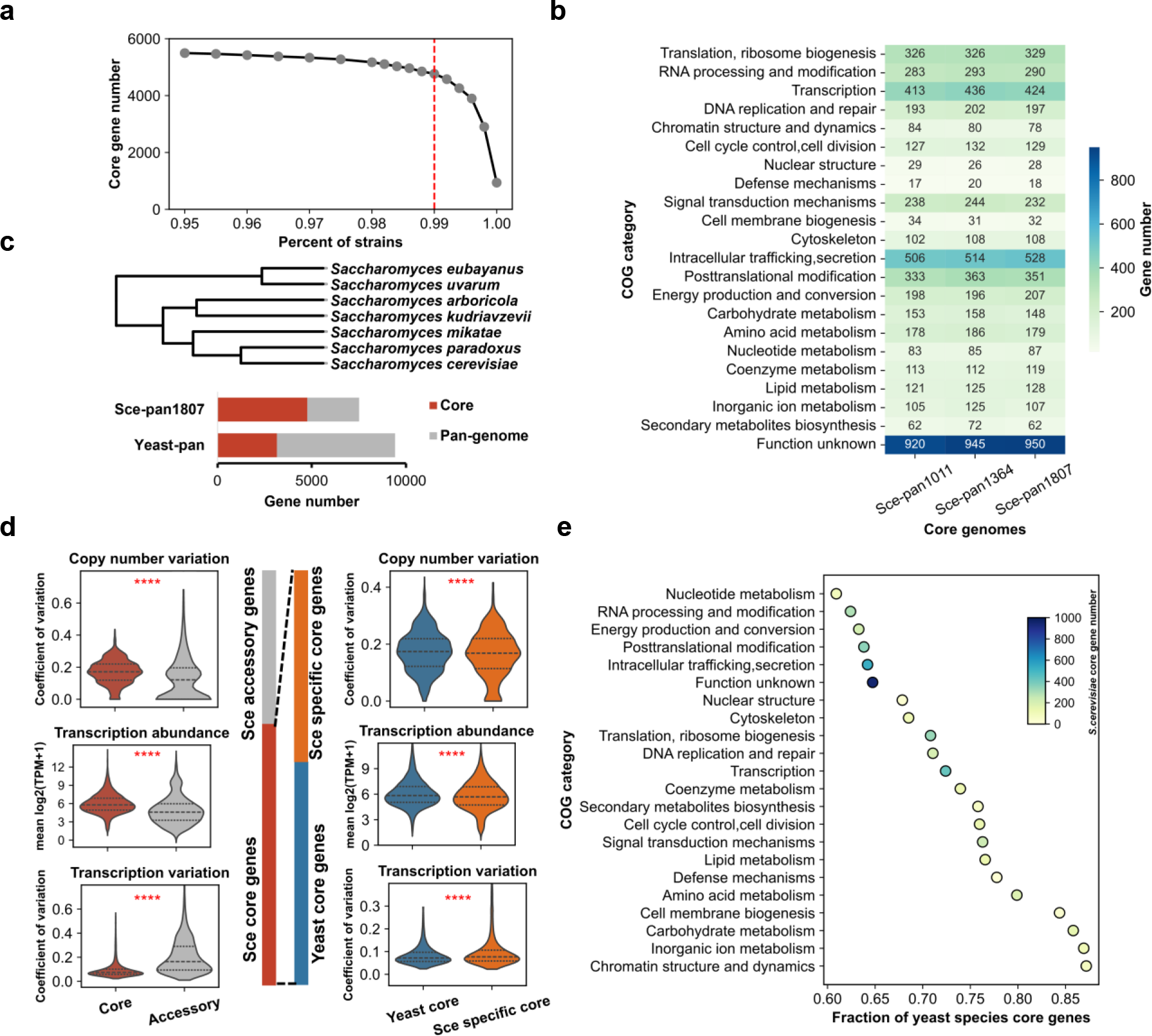
**Core genome analysis of 1,807 *S. cerevisiae* strains and its comparison with that of 7 *Saccharomyces* species**. Sensitivity analysis for core genome definition across *S. cerevisiae* isolates (a). Functional comparison of core genomes derived from three distinct pan- genomes (b). A phylogenetic relationship among 7 *Saccharomyces* species^29^, as well as the comparison of the pan-genomes between yeast species and *S. cerevisiae* (c). Comparative analysis in gene copy number, gene expression/variation between *S. cerevisiae* core and accessory genes, as well as between yeast core and non-yeast core genes. Here, the core genes from *S. cerevisiae* were further divided into yeast core and non-yeast core genes, the latter only existing in *S. cerevisiae*, according to the newly defined core genomes from 7 yeast species (d). Conservation analysis of yeast core genes across different functional categories. The X-axis represents the proportion of yeast species’ core genes in all *S. cerevisiae* core genes for each COG category (e). The statistical analysis for two group comparison was based on a two-sided T-test. Sce is an abbreviation of *S. cerevisiae.\*\*\*\**: *P* value <0.0001.

Next, the metabolic functions encoded in the 3 core genomes of *S. cerevisiae* were compared thoroughly (Fig. 2b). For each core genome, the gene functions were classified based on the COG categories. In general, the 3 core genomes exhibit remarkably similar tendencies when considering the functional distribution of core genes, highlighting the functional conservation encoded by the core genome of *S. cerevisiae*. Note that a portion of the core genes in all 3 core genomes belong to the group of unknown functions (920 for Sce-pan1011, 945 for Sce-pan1364, and 950 for this study), indicating that there exist substantial knowledge gaps in our understanding of the *S. cerevisiae* genome.

Utilizing the same approach, we assembled a pan-genome for 7 distinct *Saccharomyces* species, including *S. cerevisiae* (Fig. 2c, Supplementary Table 2). There are 9,418 ORFs in the pan- genomes of 7 yeast species, comprising 3,144 core genes and 6,274 accessory genes. When compared to the pan-genome of *S. cerevisiae*, the larger pan-genome size (9,418 vs 7,514) and the smaller core genome size (3,144 vs 4,766) for those 7 yeast species showcased the high quality of the pan- and core-genome constructed in this study.

### Evolution of the core genome within and across yeast species

Through sequence alignment between the core genomes from *S. cerevisiae* and 7 yeast species, the genes inside *S. cerevisiae* core genome can be further grouped as yeast core genes (common in 7 yeast species) and *S. cerevisiae* specific core genes (only found in a portion of 7 yeast species, Fig. 2d). It shows that the average gene copy number differed relatively slightly amongst *S. cerevisiae* strains (mean gene copy number: 1.05 for core genome versus 1.09 for accessory genome, *P* value = 3.08×10^-6^, Supplementary Fig. 2a). However, the gene copy number of core genes varied more than that of the accessory genome (mean coefficient of variation: 0.17 for core genome versus 0.13 for accessory genome, *P* value = 2.10×10^-28^, Fig. 2d). In line with these findings, the comparison between yeast core and *S. cerevisiae* specific core genes revealed a similar pattern, with yeast core genes exhibiting a marginally higher level of variation in gene copy number compared to *S. cerevisiae* specific core genes (mean coefficient of variation: 0.17 for core genome versus 0.16 for accessory genome, *P* value = 7.43 ×10^-5^, Fig. 2d). These findings highlight the subtle changes in gene copy number and variability between the core and accessory genomes, as well as between the yeast core and *S. cerevisiae* specific core genes.

Using a similar strategy, the core genes of the 7 yeast species were annotated. In each COG, the fraction of yeast core genes per *S. cerevisiae* core genes was calculated, respectively. The higher the faction of yeast core genes, the greater conservation in evolution of metabolism from specific COGs. The top 5 most conserved COGs are Chromatin structure and dynamic, Inorganic ion metabolism, Carbohydrate metabolism, Cell membrane biogenesis and Amino acid biogenesis (Fig.2e), with 4 of them involved in enzyme-driven cellular metabolism. However, some metabolism-related categories show a relatively lower degree of conservation, such as Nucleotide metabolism, RNA processing and modification, and Energy production and conversion, indicating the loss of *S. cerevisiae* core genes in other yeast species. Collectively, those differences in metabolism conservation during the evolution of the yeast core genome may contribute to phenotypic variety across yeast species^29–31^.

To explore the differences in gene expression patterns, we gathered large-scale transcriptome data of 969 *S. cerevisiae* strains from a recent study^18^, the majority of which were included in our pan-genome analysis. The comparative analysis of gene expression profiles between *S. cerevisiae* core and accessory genome revealed that core genes are more abundant but have less variance than accessory genes (*P* value = 2.14×10^-10^, Fig. 2d), which is consistent with previous study^18^. A parallel comparison between yeast core and *S. cerevisiae* specific core genes displayed a subtle but consistent trend (mean relative gene expression abundance: 6.07 for yeast core genes versus 5.81 for *S. cerevisiae* specific core genes, *P* value = 1.06×10^-7^; mean coefficient of variation: 0.086 for yeast core genes versus 0.099 for *S. cerevisiae* specific core genes, *P* value = 1.70×10^-10^, Fig. 2d). Overall, our findings show that core genes from the *S. cerevisiae* pan-genome are prone to being highly and stably expressed across strains.

### Pan-genome infer strain-specific-GEMs

To further investigate the metabolic diversity of *S. cerevisiae* isolates, the GEMs for each strain was reconstructed based on Sce-pan1807 (Fig. 3a). Initially, the pan-GEMs of *S. cerevisiae* (pan-GEMs-1011) from our previous work was updated to reflect the corrected pan-genome (Supplementary Fig. 3a)^32^. The updated pan-GEMs encompassed 4,027 reactions, 1,236 genes and 2,766 metabolites (Fig. 3b), which was named as pan-GEMs-1807. The reduction in genes in pan-GEMs-1807 compared to pan-GEMs-1011 is mostly due to the removal of ORF redundancies in the current version of pan-genome. Then, using the pan-GEMs-1807 and gene presence matrix, we reconstructed 1,807 strain-specific GEMs (ssGEMs) via an automated pipeline. The reaction sizes among these models varied from 3,794 to 4,025, with the number of genes ranging from 1,053 to 1,186. Furthermore, strains from different niches exhibited distinct distributions in metabolic network size (Fig. 3c). For instance, strains used for bioethanol production had a higher number of genes and reactions, whereas strains for dairy manifested a smaller size. A positive correlation was observed between metabolic gene number and reaction number (Pearson’s correlation coefficient =0.64, *P* value = 2.8×10^-208^, Supplementary Fig. 3b). According to our predictive analysis, 85% of the ssGEMs could simulate the strain growth under glucose minimal media conditions. However, the theoretical maximum biomass yields did not differ significantly between strains from different habitats (Supplementary Fig. 3c, 3d). The existence of a few non-viable strains may be related to auxotrophs caused by long-term adaptation or domestication. The majority of the non-viable ssGEMs could successfully predict the growth after the gap-filling step. Interestingly, we found that autotrophies predicted by the model were mainly due to the loss of crucial metabolic genes and the corresponding reactions in pathways important for precursor biosynthesis (Supplementary Fig. 3c). For example, the reaction r_1838, catalyzed by homocitrate synthase (HCS), which is essential for lysine biosynthesis^33,34^, has been found to be absent in 97 ssGEMs. Note that the loss of reactions may also be owing to the incompleteness of sequenced genomes; hence, additional physiological studies in the future are essential to verify the reliability of the predicted auxotroph.

**Figure 3.**
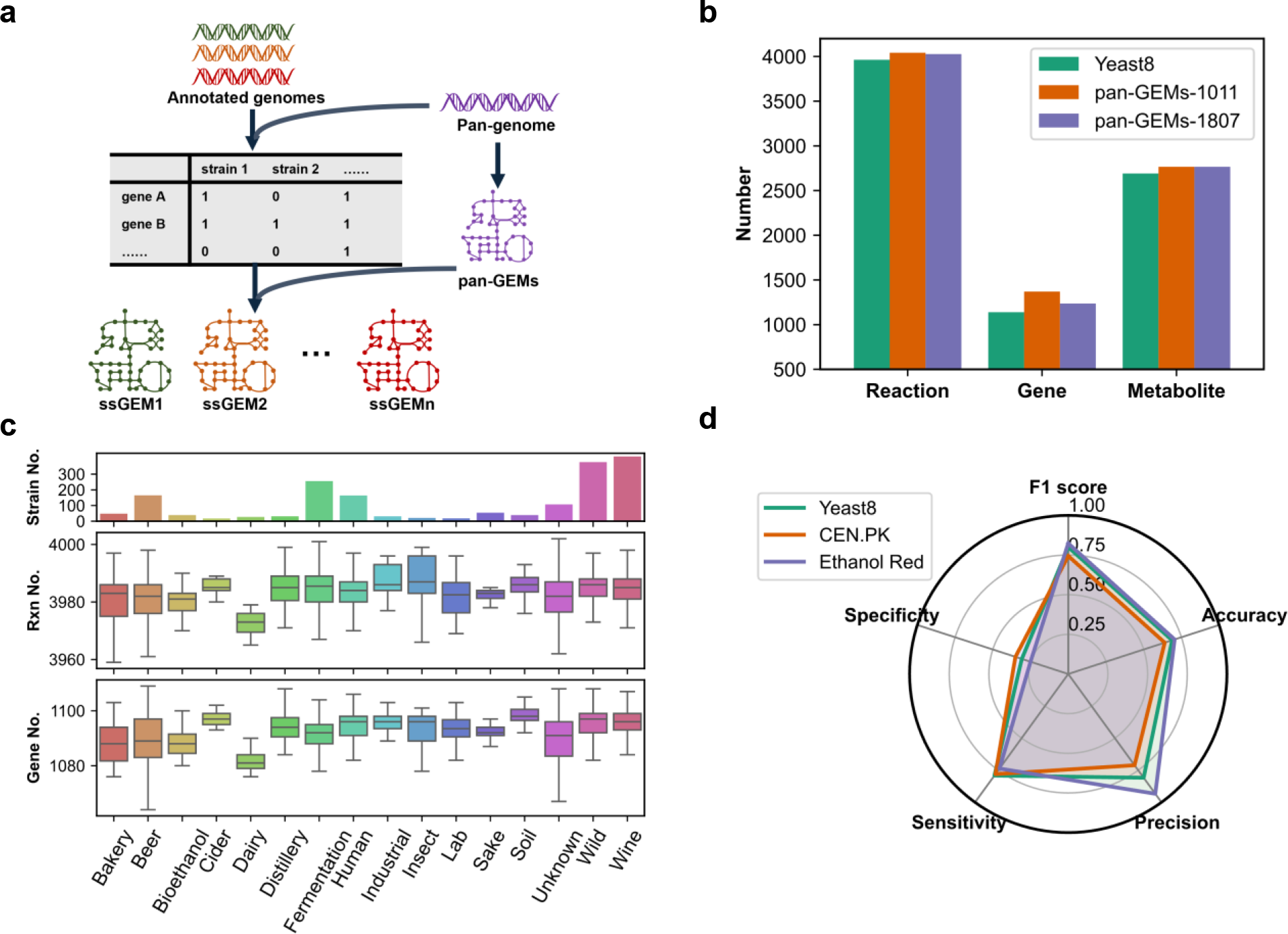
Construction and evaluation of 1,807 strain-specific GEMs (ssGEMs). Workflow for the reconstruction of ssGEMs (a). Numbers of reactions, genes, and metabolites in pan-GEMs built in this work (b). Evaluation of the number of reactions and genes in the 1,807 ssGEMs derived from different clades (c). Evaluation of the prediction capability of ssGEMs through *in silico* utilization analysis of 61 different carbon sources by three different *S. cerevisiae* strains – S288c, CEN.PK, and Ethanol red. Here the newly built ssGEMs for CEN.PK and Ethanol red were used in the comparative analysis, and Yeast8, the consesus GEMs for S288c, was used as the reference (d). The experiment dataset used for graph d is from a previous study^16^.

To further assess the quality of our ssGEMs, the utilization of 65 carbon sources by two strains- CEN.PK and Ethanol Red- was predicted, respectively, for which experimental results were already available^16^. The predictive accuracy was 0.64 for CEN.PK and 0.70 for Ethanol Red, which was comparable to that of the well-curated model -Yeast8^32^(Fig. 3d), thus lending confidence to our model prediction capability.

### Model-based analysis explore the adaptation mechanism of the transcriptional regulation of metabolic network

GEMs are widely used in omics integrative analysis^21,35,36^. To explore how the genome-wide gene expression variations shape the metabolic rewiring at a population level, the transcriptome data from 969 *S. cerevisiae* isolates under the identical culture condition with glucose as the carbon source^18^ were used to reconstruct condition-specific GEMs.

Using a well-established algorithm called GIMME^37^, we constructed 907 transcriptome-pruned ssGEMs by removing inactive reactions with low gene expression levels (Fig. 4a). Compared with the scope of the metabolic network from the clade of wild strains, we observed a reductive evolution in other clades, characterized by a smaller active metabolic network. This trend is more obvious at the reaction level than at the gene level (Fig. 4b). Examining the conservation of core reactions owned by the wild strains in other clades, some reactions were frequently lost, which may modulate the cellular metabolism (Fig. 4c, Supplementary Fig. 4a). For example, the frequently lost reaction r_0730, involving the mitochondrial formylation of Met-tRNAiMet catalyzed by mitochondrial methionyl-tRNA synthetase (MetRS), has been shown to have an incremental effect on mitochondrial translation^38,39^. Pathway enrichment analysis of the 40 most commonly lost reactions during model refining reveals that the majority of those reactions are related to cell permeability and sensitivity (Steroid biosynthesis^40^, adjusted *P* value = 4.61× 10^-8^; Glycerophospholipid metabolism^41^, adjusted *P* value = 0.01; Inositol phosphate metabolism^42^, adjusted *P* value = 0.004), cofactor metabolism (Folate metabolism^43^, adjusted *P* value = 0.02; Thiamine metabolism^44^, adjusted *P* value = 0.02), and protein synthesis (Threonylcarbamoyladenosine (t6A) metabolism, adjusted *P* value = 0.02, which is involved in a universally conserved tRNA modification^45^) (Fig. 4d).

**Figure 4.**
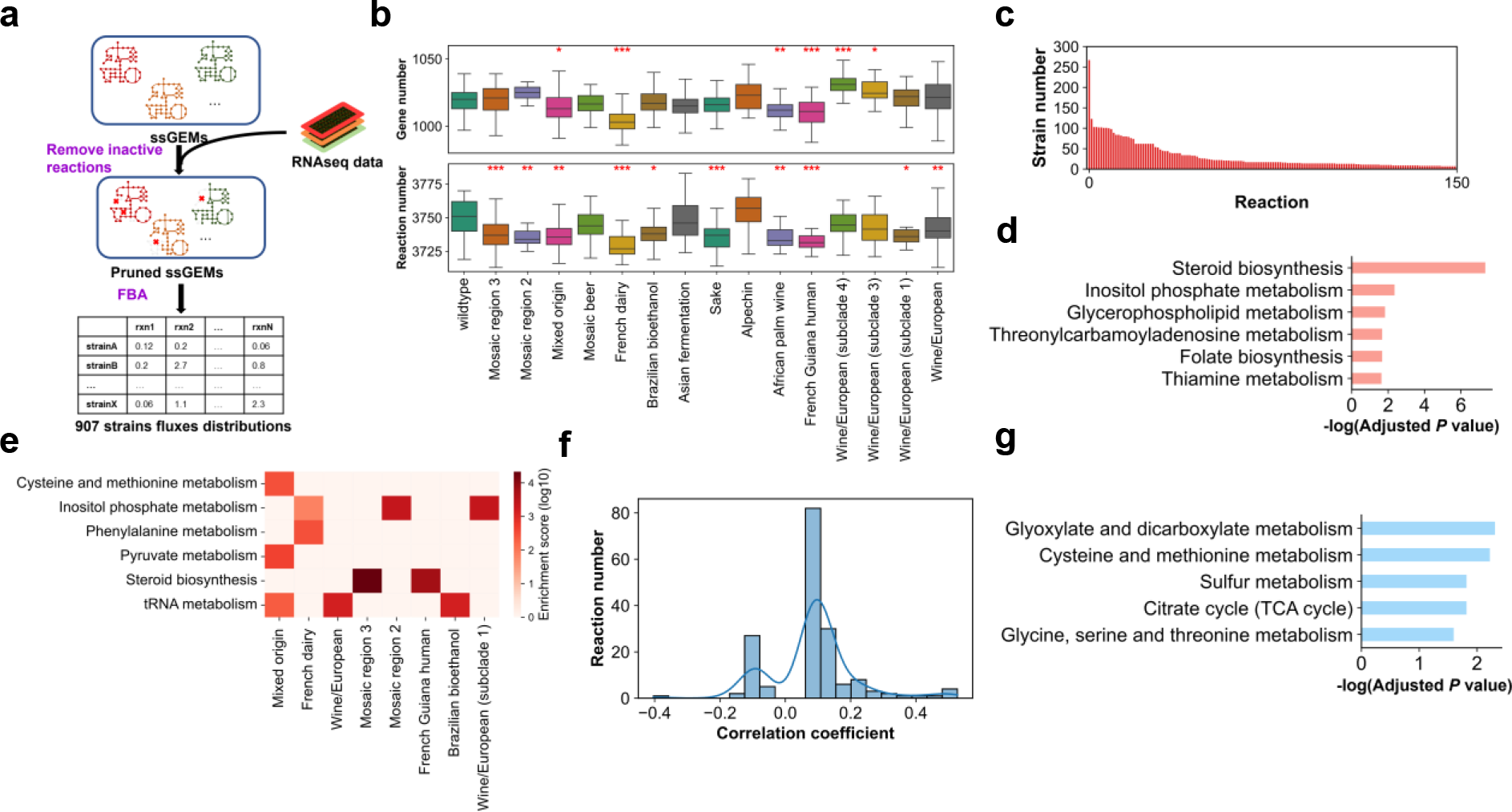
Integration analysis of ssGEMs and transcriptomics helps to explore the evolutionary mechanisms of metabolic networks. Workflow for the construction and simulation of transcriptome-pruned ssGEMs (a). Comparative analysis of metabolic network sizes across distinct evolutionary clades of *S. cerevisiae* (b). The statistical analysis of wild-type core reactions which are lost in other strains’ ssGEMs (c). Pathway enrichment analysis involving the top 40 wild-type core reactions which are lost in other strains’ ssGEMs (d). Pathway enrichment analysis of wild-type core reactions which are lost in other strains’ ssGEMs across different clades (e). Correlation analysis between metabolic fluxes and associated gene expression levels. The correlation coefficient was computed using Spearman’s rank correlation method. Only reactions with *P* values < 0.05 were considered, while reactions with *P* values > 0.05 were excluded from subsequent analysis. (f). Pathway enrichment analysis for reactions showing high correlation (g). The statistical analysis for two group comparison was based on a two-sided T-test. *: *P* value < 0.05; **: *P* value < 0.01; ***: *P* value < 0.001.

Furthermore, our findings revealed distinct patterns in reaction loss among different clades (Fig. 4e). For example, inositol phosphate metabolism, which plays an important role in membrane synthesis^42^ and stress responses^46^, experienced significant reaction loss in domesticated clades such as French dairy and Wine/European (subclade 1), which may be related to the adaptation to high osmotic stress conditions during domestication. As another example, the steroid biosynthesis, oxygen-dependent in *S. cerevisiae*^47^, is consistently inactive in strains from French Guiana human clade^48^, which mainly grow under low-oxygen conditions.

We then used the pruned ssGEMs to calculate fluxes and explore the correlation between metabolic flux and gene expression levels. Based on the measured relative growth data for 969 *S. cerevisiae* isolates^18^, we fixed the specific growth rates in the model accordingly and calculated the flux distributions by minimizing glucose uptake via flux balance analysis (FBA). After that, the correlation analysis between fluxes and gene expression levels was conducted, yielding correlation coefficients ranging from -0.40 to 0.53 (Fig. 4f). The presence of negative correlations in some reactions could be attributed to local flux coordinations^49^. Reactions were ranked based on their correlation coefficients. The top 10 reactions, which are predominantly regulated at the transcriptional level, primarily participate in specific sub-pathways, i.e., amino acid metabolism and central carbon metabolism (Supplementary Fig. 4c). Meanwhile, pathway enrichment analysis revealed that sub-pathways in amino acid metabolism (e.g., cysteine and methionine metabolism, adjusted *P* value = 0.001; glycine, serine, and threonine metabolism, adjusted *P* value = 0.01) and central carbon metabolism (like the TCA cycle, adjusted *P* value = 0.01) have stronger correlations between flux and transcription (Fig. 4g). Our findings are consistent with previous studies showing that the TCA cycle and synthesis of amino acid fluxes are predominantly regulated by transcription^50–52^. Collectively, for the first time, our integrative analysis reveals the complex regulation of metabolic fluxes at transcriptional levels on a population scale. Our analysis suggests that fluxes and transcription exhibit a weak correlation at the genome-wide level among the strains investigated, implying that changes in gene expression do not always reflect changes in fluxes^50,53^.

### Multidimensional analysis captures potential adaptative mechanism of *S. cerevisiae* strains sampled from oxygen-limited conditions

Finally, a thorough analysis that encompassed transcriptome, pan-genome, and ssGEMs was performed to understand how *S. cerevisiae* adapts to its specific ecological niche. In this scenario, we focused on the strains belonging to the dairy, bioethanol, and human clades, all of which displayed distinct metabolic networks compared to the wild type (Fig. 4b). Furthermore, strains from these three clades typically encounter oxygen limitation in their biological habitats^48^.

Initially, we extracted the pan-genome for human, dairy, bioethanol, and wild strains, respectively, from Sce-pan1807 (Fig. 5a). It shows that the human, dairy, and bioethanol populations have a relatively smaller pan-genome size compared to the wild population, albeit with a larger number of core genes. The human clade exhibited the smallest pan-genome size and the largest core genome size, potentially attributable to the stressful growth condition within the human gut^54^. This suggests that the reductive genome evolution is widespread in strains under specific ecological environments, consistent with a recent study^55^. Moreover, the variations in the core genomes among different strain clades highlight the genetic diversity arising from adaptive evolution (Supplementary Fig. 5a). Principal Component Analysis (PCA) showed that, unlike the core genome content (Fig. 5b, Supplementary Fig. 5b and 5c), both the accessory genome and the pan-genome content can be employed to effectively classify strains from clades of human, dairy, bioethanol, and wild type based on gene presence/absence or gene copy number. Afterwards, a random forest algorithm was used to extract pivotal genetic features for classifying strains based on their accessory genome (Supplementary Fig. 5d). It shows that each clade displayed unique genetic features in the accessory genome in terms of gene presence/absence and copy number (Fig. 5c, Supplementary Fig. 5e). Although some of the top feature genes have unknown functions, annotations of all feature genes suggest that they are mainly related to stress response and cell membrane/cell wall synthesis (Supplementary Fig. 6a and 6b)^56^. Additionally, the feature genes extracted based on gene presence and copy number data are different (Supplementary Fig. 6a and 6b), which hints that the variations in gene content and gene copy number may have been both modulated in response to the environmental stress.

**Figure 5.**
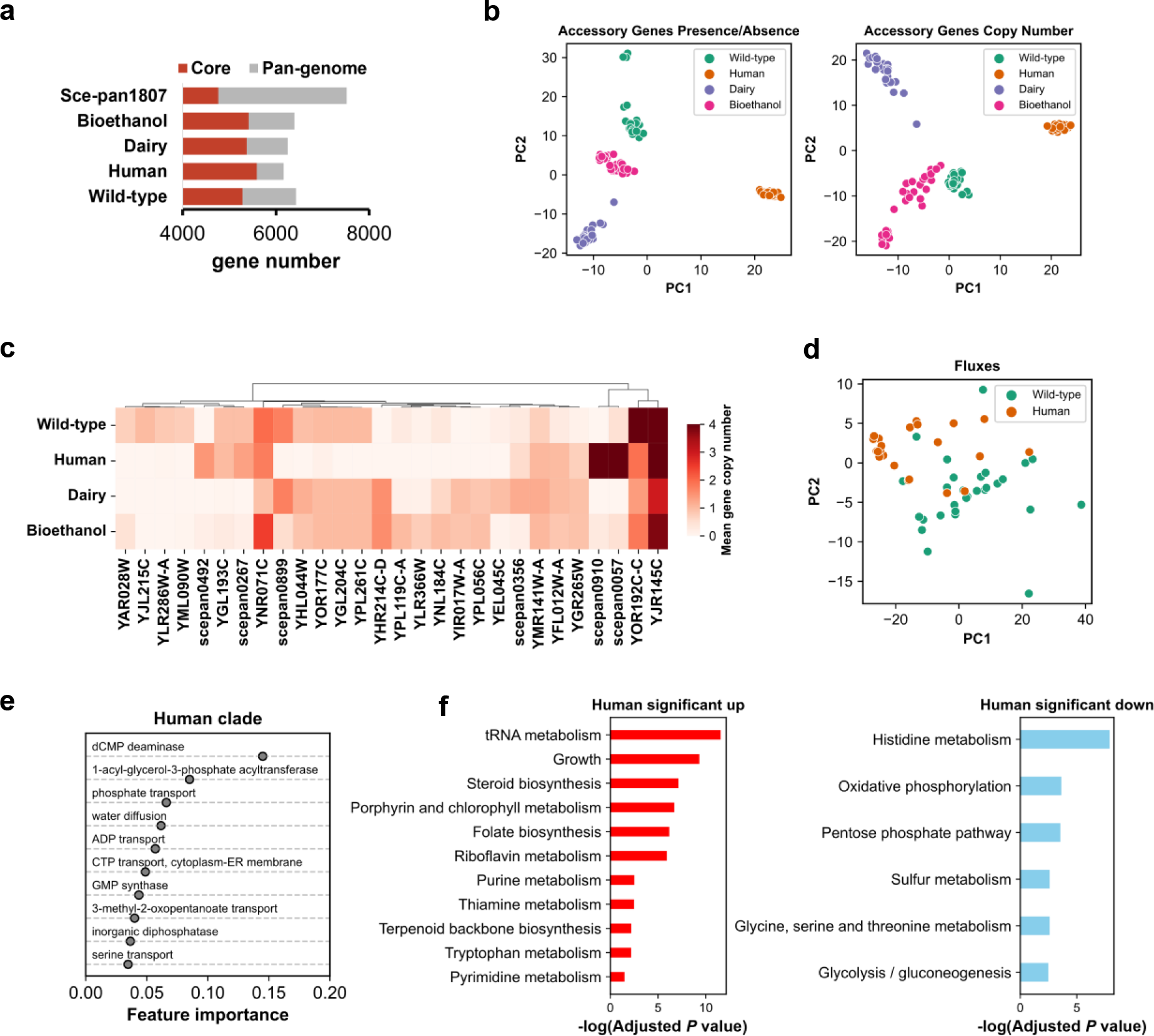
Multidimensional analysis from pan-genomes to metabolic networks helps to uncover the potential adaptative mechanisms of *S .cerevisiae* from human, dairy and bioethanol related niches. A comparison of the pan-genome among strains from the clades of bioethanol, human, and dairy and wild-type (a). PCA clustering analysis of strains from different clades based on the gene presence/absence matrix (left) and the gene copy number matrix of the accessory genome (right) (b). Top genes with copy number variation used to classify strains from different clades by the random forest algorithm (c). PCA clustering analysis of strains from the wild type and human gut environment using genome-scale fluxes obtained from ssGEMs simulation (d). Top features of metabolic fluxes (represented by the corresponding reaction names) used to classify human-related strains from the wild-type by the random forest algorithm (e). Enrichment analysis of reactions with differential metabolic fluxes between human-related strains and the wild-type (f).

Next, we investigated the metabolic flux rewiring mechanisms underlying *S. cerevisiae* strains from human, dairy, and bioethanol clades through ssGEMs simulation. To accurately capture the trends in metabolic flux variations across various strains, herein we integrated both relative growth data and transcriptomics into ssGEMs for simulation^57^, during which the relative growth rate was just set as a soft constraint (Material and Methods). The predicted growth rates are consistent with the experimental data (Pearson’s correlation coefficient = 0.85, *P* value = 3.68 ×10^-28^, Supplementary Fig. 7a). Additionally, our simulation successfully captured the phenotype of bioethanol strains, which shows higher ethanol yield than wild strains^58^. Interestingly, the human clade also exhibits a higher ethanol secretion compared to the wild strains, potentially due to its adaptive evolution in an oxygen-limited human gut environment (Supplementary Fig. 7b)^48^. Overall, the reliability of our pipeline in condition-specific flux simulation is demonstrated by the precise predictions for growth and ethanol secretion of *S. cerevisiae* strains from different clades.

We ask whether the metabolic variations among the selected strains from the above 4 different clades could be captured by flux analysis. We found that, unlike the bioethanol clade, the human and dairy clades could be clearly distinguished from the wild type strain through PCA analysis only based on flux distributions (Fig. 5d, Supplementary Fig. 7c). However, the differentiation of strains within the four clades based on metabolic fluxes did not achieve the same resolution as clustering based on gene presence or gene copy numbers (Fig. 5b, Supplementary Fig. 7c), which is likely due to the fact that all RNA-seq datasets used in our analysis were collected under the same growth condition^18^. This also hints that the metabolic fluxes, compared to the genomic variation, may be more conserved in maintaining cellular fitness. Subsequently, with the strains from the wild type clade as the reference, the key metabolic features were delineated for the human, dairy, and bioethanol clades through machine learning, respectively (Fig. 5e, Supplementary Fig. 5b, 7d, 7f). Specifically, within the human clade, dCMP deaminase was identified as a pivotal metabolic feature. The down-regulation of flux through the reaction catalyzed by dCMP deaminase might play a role in maintaining the balanced dNTP pool required for proper DNA metabolism^59^(Fig. 5a). We further evaluated the metabolic variations at the pathway level (Fig. 5f, Supplementary Fig. 7g, 7h). Interestingly, we found that there are many similar metabolic signatures among strains from the human, dairy, and bioethanol clades. For example, both human and bioethanol strains, likely adapted to oxygen-limited environments, exhibited significant suppression of the oxidative phosphorylation pathway.

Intriguingly, dairy and human strains showed nearly identical patterns in metabolic variation, with upregulated pathways predominantly associated with growth, including tRNA metabolism^60^, steroid biosynthesis^61^, terpenoid biosynthesis^62^, purine metabolism^63^, and pyrimidine metabolism^64^. This result is consistent with the observed faster growth rates of both human and dairy strains compared to wild strains^18^(Supplementary Fig. 7f). Additionally, a significant downregulation of histidine metabolism in both human and dairy clades is consistent with findings from a recent comparative metabolomics study of *S. cerevisiae*^65^ (Fig. 5f, Supplementary Fig. 7h). These shared metabolic variations among human, dairy, and bioethanol clades underscore the pervasive signatures of convergent evolution within the metabolic networks of independently adaptive subpopulations. As for strains with human clade, we discovered that the upregulation of sub-pathways involved in cofactor metabolism, particularly thiamin metabolism (*P* value = 0.003), which generates the indispensable cofactor thiamine pyrophosphate (TPP) for enzymes participating in central carbon and energy metabolism^66^, has been documented at both metabolic flux and transcriptional levels (Fig. 5f, Supplementary Fig. 8a). This upregulation may coincide with the adaptative evolution of strains inside the human gut environment, which is possibly driven by the resource competition among microbial species within the human gut^54,67^.

## Discussion

The microorganisms always undergo adaptive evolution to improve cellular fitness, whereby the genome compositions across strains from the same species are shaped by environments to promote trait diversity^9,68,69^. In this study, we recreated a new-version of pan-genome for *S. cerevisiae* (Sce-pan1807) using large-scale genome datasets and evaluated its reliability in reflecting genomic and phenotypic evolution across 1,807 distinct strains. The Sce-pan1807 encompasses 7,514 distinct gene families, comprising 4,766 core genes and 2,748 accessory genes. Comparison with pan-genomes from earlier studies^9,20^ revealed the higher comprehensiveness of our pan-genome owing to the delicate workflow and massive genome dataset used in this work (Fig. 1b, c). Note that, even for single species, the size of the *S. cerevisiae* pan/core-genome can vary throughout each iterative upgrade due to differences in gene cluster algorithms, the number of sampled strains, and the definition of pan/core genes^9,20,70^. Given that a fair number of new genes, which not existing in model strains - *S. cerevisiae* S288c/CEN.PK 113.7D, were identified through the pan-genome analysis, the extra molecular experiment to verify the functions of those additional genes will certainly deepen our understanding of how the gain of genes impacts cellular fitness. Currently, our pan-genome analysis is primarily focused on gene existence/copy analysis across diverse strains. In further comparative genomic analysis, the structural variants (SVs) and single nucleotide polymorphisms (SNPs), which widely exist in yeast genomes^9,71^, should also be taken into account to illustrate how multidimensional evolutionary signatures contribute to trait diversity.

The strain phenotypes are actually determined by the interaction of multilayer molecular networks within cells^72^. As one of the most important computational tools for systems biology studies, genome-scale metabolic models (also as GEMs) could be used as the scaffolding to transform big data into knowledge^21^. Based on our Sce-pan1807, we have successfully reconstructed ssGEMs for 1,807 *S. cerevisiae*. Importantly, each ssGEMs could mimic the metabolic potential of the isolate when considering the gene gain and loss information. However, considering the huge number of ssGEMs developed in this work, more physiological datasets are required to further evaluate the prediction capability of those ssGEMs in order to apply these models in more detailed studies. Only GEMs may not accurately reflect in vivo cellular metabolic activities^72^. Here, guided by large-scale transcriptomics from a recent study^18^, ssGEMs for 907 isolates were refined. The pruned genes/reactions from ssGEMs reflected the reductive evolution in most domesticated clades, indicating human-related activities actually modulate the *S. cerevisiae* evolution. The metabolic regulation at the transcription level in *S. cerevisiae* was also explored using ssGEMs simulation. The population-scale correlation analysis reveals that a few parts of pathways, i.e., amino acid synthesis, were substantially regulated at the transcription level, thus substantiating previous knowledge obtained from studies at the single strain level^52^. Specially, to illustrate the unique advantages of ssGEMs- based pan-genome analysis, we selected human, dairy, and bioethanol clade^9^ as representative examples to explore the adaptive mechanisms of *S. cerevisiae* strains under specific ecological niches, which has never been done before. Employing an integrative analysis incorporating pan- genome, transcriptome, and ssGEMs, some interesting findings can be achieved. For example, we found that, compared to wild strains, the strains from the human gut environment exhibit remarkable differences at multiscale levels, including gene presence/absence, gene copy number, and fluxes of the metabolic network, all of which may contribute to the adaptation underlying the human gut environment with limited oxygen supply and intense resource competition. Thus, our analysis here provides a universal paradigm for profiling the metabolic rewiring of *S. cerevisiae* at a holistic level, which could be applied to examine the long-term evolution of other valuable microorganisms.

In conclusion, we have recreated the new-version of pan-genome and large-scale strain-specific genome-scale models for 1,807 strains of *S. cerevisiae*. These resources together systematically illustrate how *S. cerevisiae* strains flexibly adapt to various stressful conditions through multidimensional evolution at both genomic and metabolic levels. As additional omics and phenotypic data become available, these yeast’s computational resources can be further updated and eventually accelerate the systematic evolutionary studies of yeast species.

## Materials and Methods

### Genome sequences collection and annotation

Totally, 1913 assembled genomes for *S. cerevisiae* were collected from published articles^9,11,13,14,16,17,20^ and the National Center for Biotechnology Information (NCBI) database. Initially, repetitive elements and low complexity DNA sequences within the genomes were identified and soft masked utilizing RepeatMasker v4.0.7 (parameters: “-species ’*Saccharomyces cerevisiae*’ -xsmall -e ncbi”, http://repeatmasker.org). This step was crucial to minimize the interference from these sequences in downstream analyses. Subsequently, gene prediction was performed using two distinct annotation tools: MAKER2 ^73^ and liftOver^74^ (https://github.com/wurmlab/flo) were employed to predict genes for each assembled genomes. The MAKER2 annotation pipeline incorporated the ab initio gene prediction method AUGUSTUS, fine-tuned with *S. cerevisiae* specific parameters (-- species=*saccharomyces_cerevisiae*_S288c). Concurrently, a homology-based approach was employed through liftOver, utilizing all transcripts of the S288c strain (http://sgd-archive.yeastgenome.org/sequence/S288C_reference/genome_releases/) as a reference for gene prediction. Finally, the gene prediction results obtained from the above methods were combined using customized scripts based on the following criteria: (1) sequences from either LiftOver or MAKER shorter than 30 residues were discarded; (2) sequences from LiftOver containing an internal stop codon were also discarded; (3) If overlapping genes exist, genes from LiftOver were used.

### Genome assessment

To ensure the high quality of genome datasets utilized for pan-genome reconstruction, a quality control process adapted from the previous study^20^ was implemented. Initially, genomes with suboptimal assemblies were excluded based on the N50 value and assembly level. The N50 values of genomes were calculated using a customized script. Genomes exhibiting contig-level assemblies or N50 values below 10kb were excluded. Next, the completeness of all annotated proteomes and assemblies was assessed using the *Saccharomycetes*_odb10 database through the application of BUSCO^24^. Genomes with assembly completeness scores or proteome completeness scores below 95% were eliminated. Afterwards, a total of 1,807 proteomes were retained for subsequent pan-genome construction.

### Pan-genome reconstruction of *S. cerevisiae*

For the construction of the *S. cerevisiae* pan-genome, a customized reference-based pipeline was employed. Initially, the *S. cerevisiae* S288c proteome was chosen as the reference owing to its comprehensive and detailed annotations by the yeast community (http://sgd-archive.yeastgenome.org/sequence/S288C_reference/genome_releases/). Using BLASTp, all proteomes were aligned against the reference to extract the non-reference genes by MMseqs2. A query gene could be grouped as the reference gene if the corresponding Percentage Identity (PID) is over 70% and the Coverage (COV) over 50% during the cluster analysis. Genes exhibiting notably high PID but limited COV scores (PID > 95%, COV < 50%, E value < 0.00001) were identified as incomplete reference genes, which were excluded from further analysis. Additionally, all proteins from the CDH.re strain were excluded due to the potential prokaryotic contamination. This preprocess resulted in a total of 104681 non-reference genes, averaging 71 non-reference genes per genome and ranging from 0 to 556 across strains. Subsequently, incomplete genes containing ’X’ in their protein sequences were systematically eliminated, resulting in 92977 non-reference genes. These non-reference genes were then integrated with the S288c proteome to undergo sequence clustering via MMseqs2’s easy-cluster function (-c 0.5 --min-seq-id 0.7 -e 0.00001 --cov-mode 0 --cluster-mode 0), yielding 7,551 ORFs.

To optimize the pan-genome for enhanced functional annotation, a specific set of criteria was established to select representative sequences from gene clusters: (1) Within a cluster, if only one sequence belongs to the S288c strain, that S288c sequence is chosen as the representative sequence. (2) Within a cluster with multiple sequences from the S288c strain, priority is given to genes already present in Yeast8^32^. If not present in Yeast8, the longest S288c sequence serves as the representative sequence. (3) In the absence of any S288c sequence, the original representative sequence from MMseqs2 is selected. Subsequently, an all-vs-all BLAST was conducted on all representative sequences of the ORFs using MMseqs2’s easy-search function. This process led to the removal of 13 incomplete genes (PID > 95%, query COV < 50%, and target COV > 90%) and 24 similar genes (PID > 70%, COV > 50%), resulting in a final set of 7514 representative genes. The functional annotation of the pan-genome was conducted utilizing the eggNOG-mapper online service with default parameters^75^(http://eggnog-mapper.embl.de/).

Lastly, the presence/absence information and copy number of each gene encompassed in the pan-genome across all strains were acquired by performing BLASTp for the 7,514 pan-genome representative genes against each strain’s proteomes with the unified threshold: PID > 70%, COV > 50%, and E value < 0.00001. Based on the gene presence/absence content analysis, a sensitivity analysis of the core genome size was performed, considering the frequency of open reading frames (ORFs) occurring in different strains. The percentage threshold defining the core genes was tested, ranging from 95% to 100%. Ultimately, genes present in more than 99% of the 1,807 strains included in the study were defined as the core genes of *S. cerevisiae*.

### Pan-genome reconstruction of 7 different yeast species

In the reconstruction of pan-genome for yeast species evolutionarily close to *S. cerevisiae*, genomes from another six yeast species closely related to *S. cerevisiae* were utilized. These species included *Saccharomyces arboricola, Saccharomyces eubayanus, Saccharomyces kudriavzevii, Saccharomyces mikatae, Saccharomyces paradoxus, and Saccharomyces uvarum*. The genomic data for these species were sourced from a prior study^29^. For the reference genome, the genome of *S. cerevisiae* S288c was selected. The reconstruction of the pan-genome for these yeast species followed the same pipeline as previously described for the *S. cerevisiae* pan- genome. This process resulted in a pan-genome for yeast species comprising 9,418 ORFs.

### ssGEMs reconstruction

In order to reconstruct a pan-GEMs for *S. cerevisiae*, the pan-GEMs^32^ of 1011 *S. cerevisiae* strains from a previous study was used as the draft template model for this work. The newly acquired pan-genome, Sce-pan1807, was aligned with the prior Sce-pan1011, employing a bi- directional best hits (BBH) alignment strategy to map the new panID to the old panID, in which the parameters were set as: PID > 70%, COV > 50%, and E value < 0.00001. According to the alignment result, we updated the pan-GEMs accordingly for 1,807 *S. cerevisiae* isolates: (1) 82 new pan-genes were incorporated, leading to the modification of 89 reactions with updated gene-protein-reaction (GPR) associations, without introducing new reactions. (2) 221 genes were eliminated from the pan-GEMs that were not found in the new pan-genome, Sce-pan1807, likely attributable to redundancies in the previous pan-genome. In total, the update involved the modification of 303 genes and the adjustment of 639 reactions’ GPR associations in the new version of *S. cerevisiae* pan-GEMs.

Subsequently, strain-specific GEMs (ssGEMs) were reconstructed for 1,807 strains, utilizing the updated pan-GEMs and the gene presence/absence matrix. A Python function was developed to generate ssGEMs automatically according to the following rules: (1) All absent genes were systematically excised from the template model. (2) Guided by the GPR associations, if all associated genes were absent, the corresponding reaction was also eliminated. (3) If any subunit of a complex was absent, the corresponding reaction was removed. (4) If only one gene from a set of isoenzymes was lost, the corresponding reaction was kept.

### ssGEMs simulation of different substrate utilizing

To evaluate ssGEMs quality, the utilization of 65 carbon sources by 3 representative *S. cerevisiae* strains-S288c, CEN.PK and Ethanol Red^16^, was collected. To simulate the utilization of a substrate in silico, the ssGEMs were constrained by the minimal media condition, in which the standard carbon source, that is glucose, was substituted by the target substrate. The simulation of growth on diverse carbon source was executed by setting the exchange rate of the relevant substrate at 1.0 mmol/gDW.h. The threshold of all simulated growth rates was set at 0.01 h^−1^ to represent the normal cellular growth on these substrates. The performance metrics applied to evaluate the simulations were defined as follows:

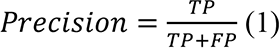

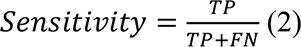

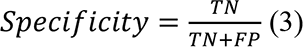

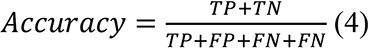

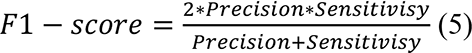

Where TP represents the number of correctly predicted cases of growth on a given substrate, FP represents the number of incorrectly predicted cases of growth on a given substrate, FN represents the number of incorrectly predicted cases of non-growth on a given substrate, and TN represents the number of correctly predicted cases of non-growth on a given substrate.

### Transcriptome-pruned ssGEMs construction and simulation

The transcriptomics of 969 *S. cerevisiae* isolates under identical culture conditions were collected from a recent study^18^. Based on the strain ID, 907 strains matched to our ssGEMs. The GIMME algorithm, as implemented in the COBRA Toolbox^37^, was utilized to generate transcriptome-pruned species-specific Genome-scale Metabolic Models (ssGEMs). For each strain, the expression profile was ranked from highest to lowest according to abundance. The reaction expression score threshold was set at the abundance value in the 75th percentile position, according to the method described in a previous study^76^. However, as the GIMME algorithm only removes inactive reactions from models, a customized Python script was implemented to remove all unused genes lacking corresponding reactions from the GIMME output models, resulting in 907 refined ssGEMs.

To simulate the flux distributions using the refined ssGEMs, the following steps were employed:

1. Collecting growth data: The growth times reaching the mid-log phase (t-mid time) for each strain were collected from the batch cultivation^18^ and converted into relative growth rates by taking the reciprocal of the t-mid time.
2. Relative growth rate calculation: Then, the top 5% fastest strains were selected as the reference, and each strain’s relative growth rate was calculated by comparing it to the reference. All values greater than 1 were set to 1, ensuring that all values ranged from 0 to 1.
3. Simulation: According to a previous study^77^, the maximum growth rate was set at 0.48 h^-1^. Based on the above relative growth rate and the maximum growth rate, the corresponding specific growth rate for each strain was re-scaled. The growth rate in the model was fixed, and glucose uptake was minimized via FBA in COBRApy^78^ to obtain the flux distributions for each strain.

To quantify the correlation between metabolic flux and gene expression level, the Spearman correlation coefficient method was employed using the SciPy (https://scipy.org/) package in Python. Pathway enrichment analysis for specific reaction sets, such as reactions with significant correlation between expression and metabolic fluxes or common lost reactions among all strains, was conducted using the enrichr module in the GSEApy^79^ package. For the parameters used as input in the GSEApy package function, reference reaction sets used for pathway enrichment analysis were extracted from the Yeast8 model^32^, with the adjusted *P* value cutoff set at 0.05.

### Pan-genome analysis for strains from specific evolutionary clades

The reconstruction of the pan-genome for distinct clades, including but not limited to wild-type, human, bioethanol and dairy clades, was based on the pan-genome Sce-pan1807. As for the clade-specific pan-genome, any pan-gene absent from all strains within a given clade was excluded. Genes that present in all strains within a given clade were defined as core genes.

### Model simulation for strains from clade of bioethanol, dairy, human and wild-type

Transcriptome integration and model simulation were conducted using RIPTiDe (v3.4.81), in a manner similar to that described in a previous study^80^. Steps are as follows:

1. Preparing transcriptome. The transcriptome from ^18^ for each strain was prepared in in Transcripts Per Million (TPM) format.
2. Setting the known ethanol yield as the lower bound. This step was specially designed to overcome the limitations of classical GEMs, which cannot capture the metabolic overflow phenomenon existing in Crabtree positive yeasts. A lower bound of 0.18 g/g glucose was set for the ethanol yield, based on the lowest ethanol yield of *S. cerevisiae* measured in batch culture under different growth conditions^81^.
3. Setting the relative growth rate as a kind of soft constraint. Maximization of growth was set as the objective for all ssGEMs. The relative growth rates derived from the method described in the “Transcriptome-pruned ssGEMs construction and simulation” section were set as the lower bound of the fraction of growth rate to the maximal growth rate for each strain through the “fraction” parameter in the contextualize() function of RIPTiDe^57^.
4. Flux simulation. The contextualize() method in RIPTiDe was executed to compute the flux distributions for each strain. For each strain, the final flux distribution was determined by averaging all sampled fluxes, which were then utilized for subsequent analyses.

### Explore potential genetic/metabolic features with machine learning

To evaluate the genome-scale genetic/metabolic differences among the different clades, the PCA clustering algorithm was performed using gene presence/absence content, gene copy number content, or metabolic flux profiles as input. The PCA analysis was implemented using the sklearn.decomposition.PCA module in the scikit-learn package(https://scikit-learn.org/stable/).

Next, to extract the potential genetic/metabolic features for each clade, the Random Forest classification algorithm was implemented using the scikit-learn package(https://scikit-learn.org/stable/). Gene presence/absence content, gene copy number content, or metabolic flux profiles, labeled with clade type were respectively set as input. The entire dataset was split into five folds. In each iteration, four folds were used to train the model with optimized hyperparameters, and the trained model was tested on the remaining fold to calculate the accuracy score. This step was repeated five times to ensure that each fold was used as a test dataset. The average accuracy of those five test accuracies was used. All other parameters were set to their default values. The average feature importance from the best estimator in each fold of the 5-fold cross-validation process was used to score and rank the genetic/metabolic features.

### Differential expression and KEGG pathway enrichment analyses

Clade-specific gene differential expression analyses were performed by DESeq2 method^82^ in OmicVerse package^83^. The thresholds to identify significantly expressed genes were established at adjusted *P* value < 0.05 and absolute log2 fold-change > 2. KEGG pathway enrichment was performed by enrichr module in GSEApy package^79^. Reference gene sets were extracted from the KEGG database^84^, with the adjusted *P* value cutoff set at 0.05.

### Statistical analysis

For two group comparisons in this work, two tailed Wilcoxon rank sum tests were performed using the SciPy (https://scipy.org/) package in Python.

### Data availability

All scripts are accessible at https://github.com/hongzhonglu/Unified_Yeast_GEMs_Database. All large files, including ssGEMs, annotated genome data, and transcriptome-constrained ssGEMs are accessible at https://figshare.com/s/9c2faecc9d79d4825d0d.

## Supporting information

S. cerevisiae strains information

7 yeast species genomic information

## Acknowledgements

This work is supported by grant 2022YFA0913000 from the National Key R&D Program of China, Shanghai Pujiang Program, and grant 22208211 and 22378263 from the National Natural Science Foundation of China (NSFC). The funding body has no role in the design of the study, analysis and interpretation of the data, preparation of the manuscript, and decision to submit the manuscript for publication.

## Author contributions

HZL and YJZ designed the research. HYW performed the research. All authors interpreted the results, discussed, drafted and approved the final manuscript.

---

**Supplementary Figure 1.**
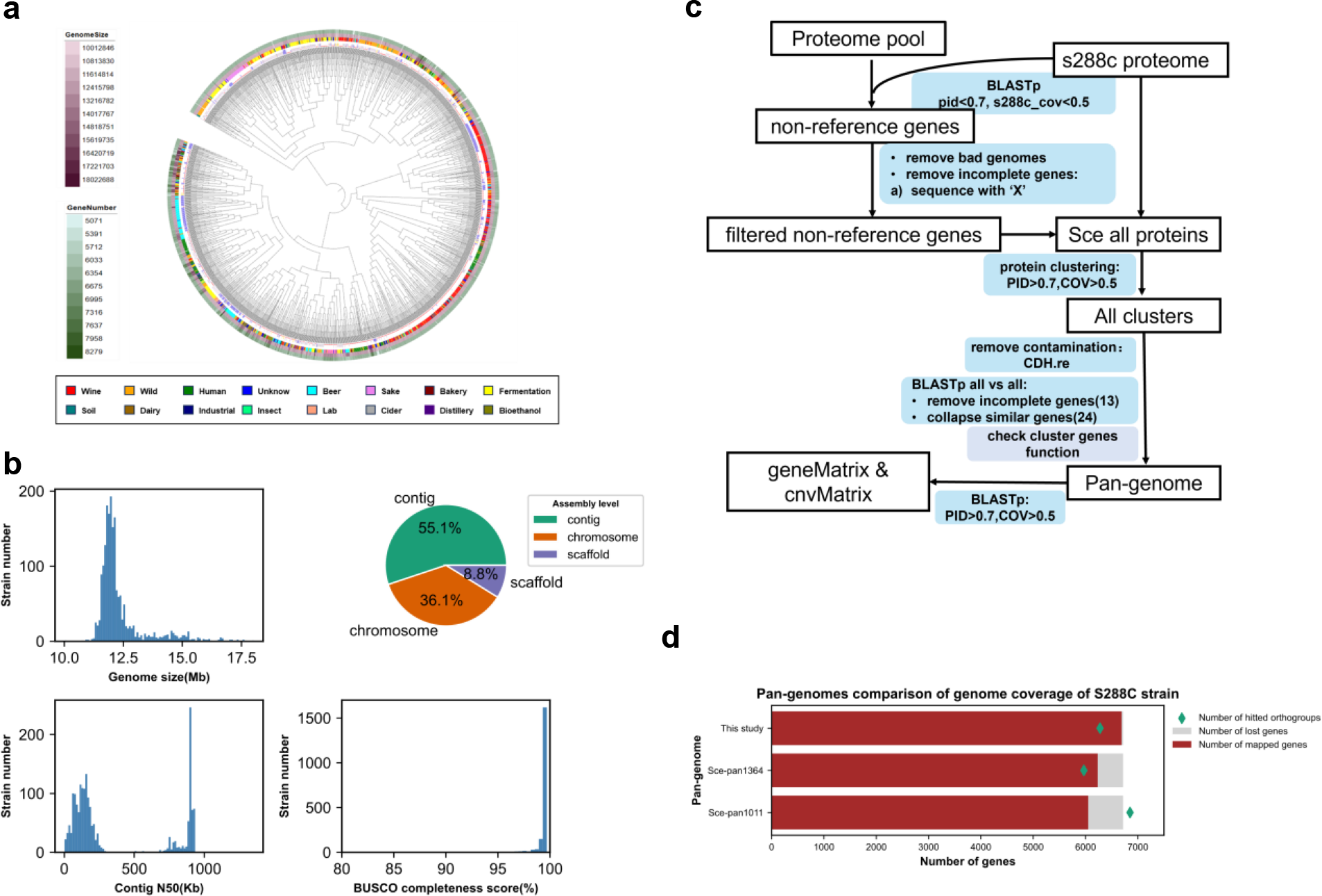
Comprehensive genome dataset of S. cerevisiae isolates and pan- genome construction. a, Unrooted neighbor-joining tree of 1913 *S. cerevisiae* isolates based on whole-genome Average Nucleotide Identity (ANI) data. The circular dendrogram is annotated with genomic characteristics of various strains, detailed from the inner to the outer rings: ecological niche classifications, genome sizes ranging from 10 to 18 Mb, and gene counts varying between 5071 and 8279. b, Statistical evaluation of genome size, assembly quality metrics (N50), and assessment of genome completeness for all isolates. c, The methodology employed for constructing the pan-genome. d, Genome coverage analysis of the reference S288C strain for 3 pan-genomes. ‘Number of hitted ORFs’ indicates the number of pan genes that have been detected in reference genome. ‘Number of lost genes’ indicates the number of reference strain’s genes that have not been detected in pan-genome. ‘Number of mapped genes’ indicates the number of reference strain’s genes that have been detected in pan-genome.

**Supplementary Figure 2.**
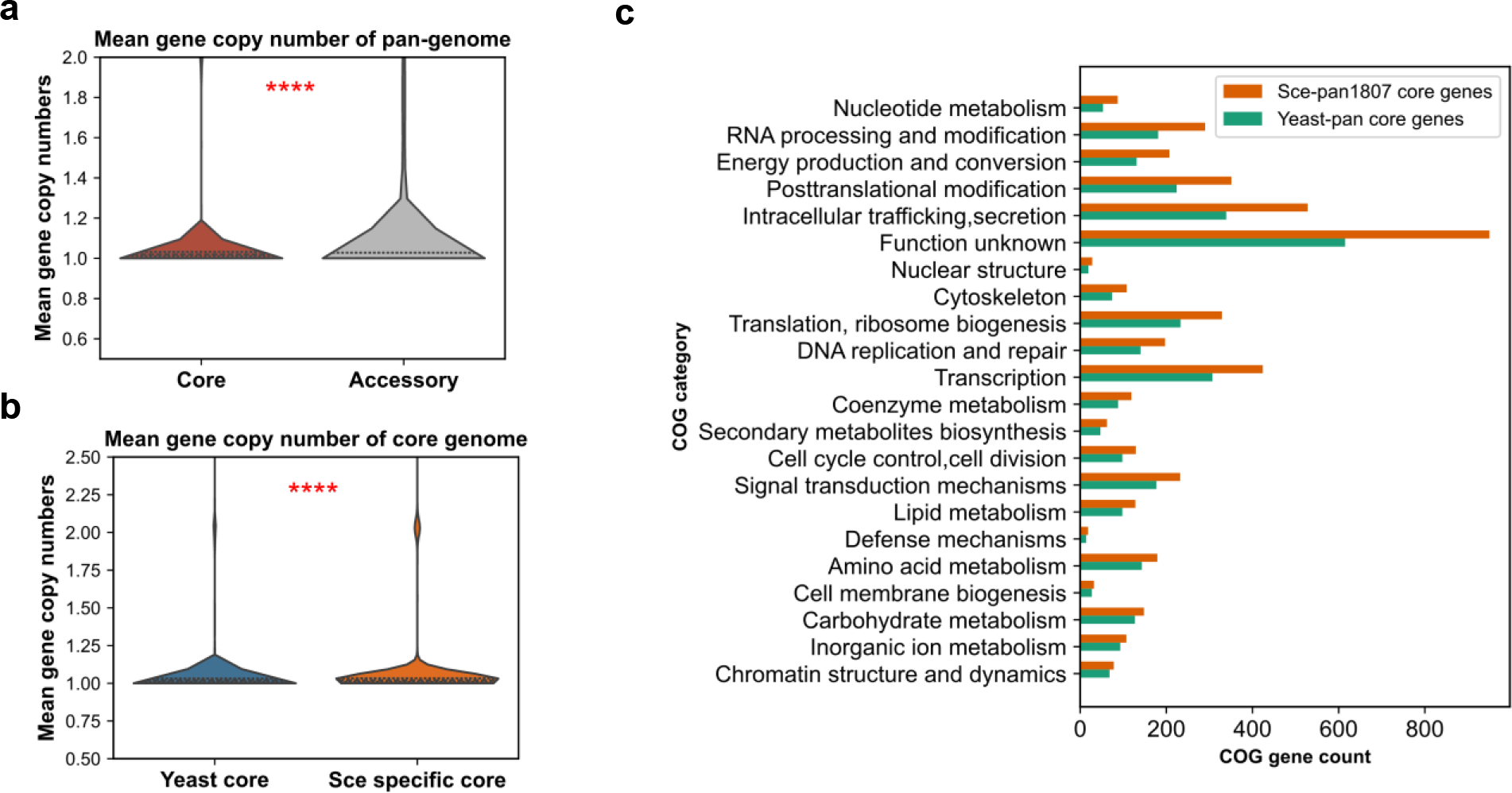
Core genome analysis of *S. cerevisiae* and yeasts species. Comparative analysis of gene copy number between core and accessory genes (a), as well as between yeast core and non-yeast core genes (b). Statistical significance was determined using a two-sided T-test. Core genome functional comparison between Sce-pan1800 and Yeast species pan-genome (c). ****: *P* value < 0.0001.

**Supplementary Figure 3.**
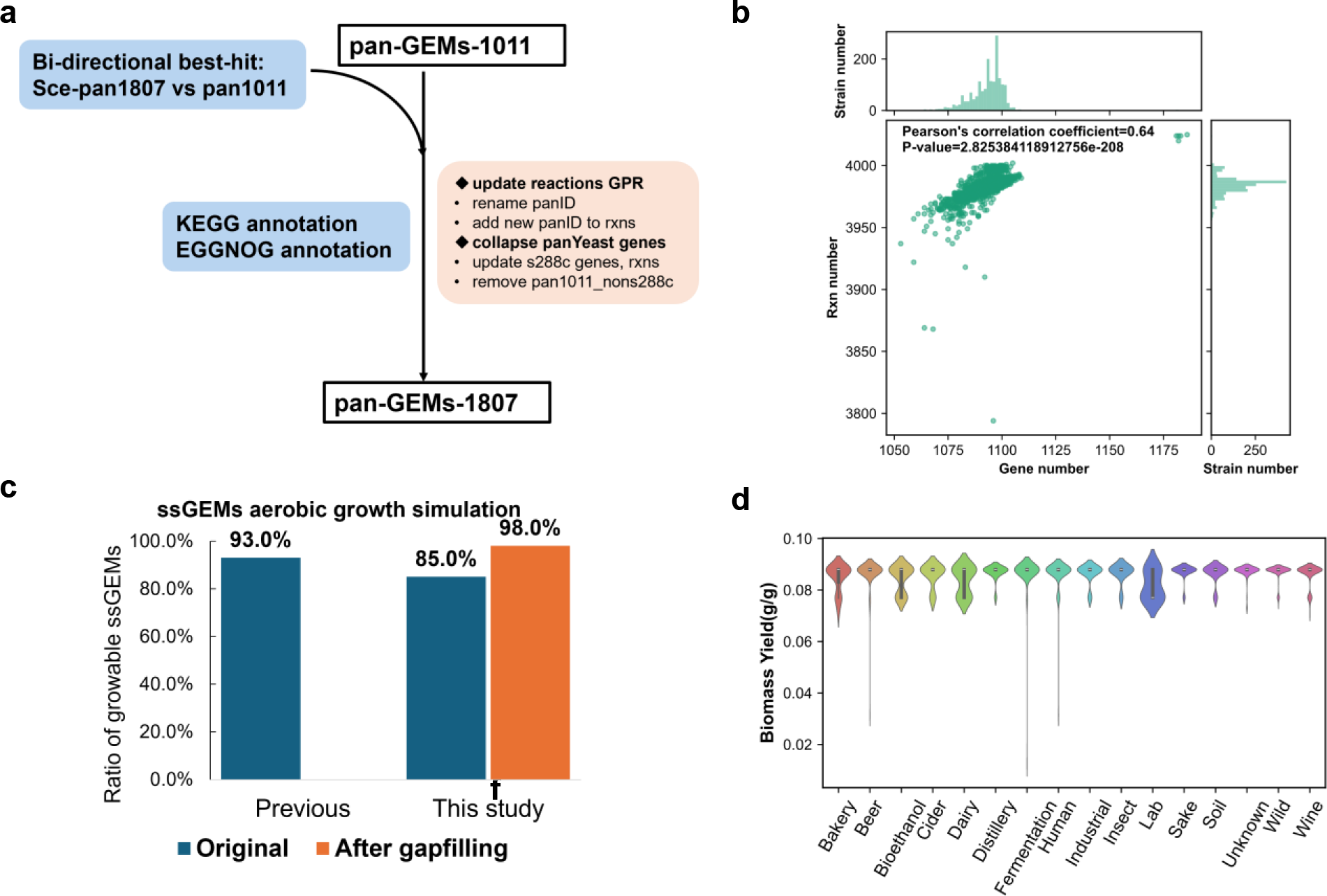
Construction and evaluation of 1807 strain-specific GEMs. The methodology employed for updating the panYeast model (a). Evaluation of the number of reactions and genes for each ssGEMs (b). In silico growth simulation results on minimal media with glucose as the sole carbon source for ssGEMs from this study compared to a previous study (c). Comparison of predicted theoretical maximum biomass yields across strains from different ecological niches (d).

**Supplementary Figure 4.**
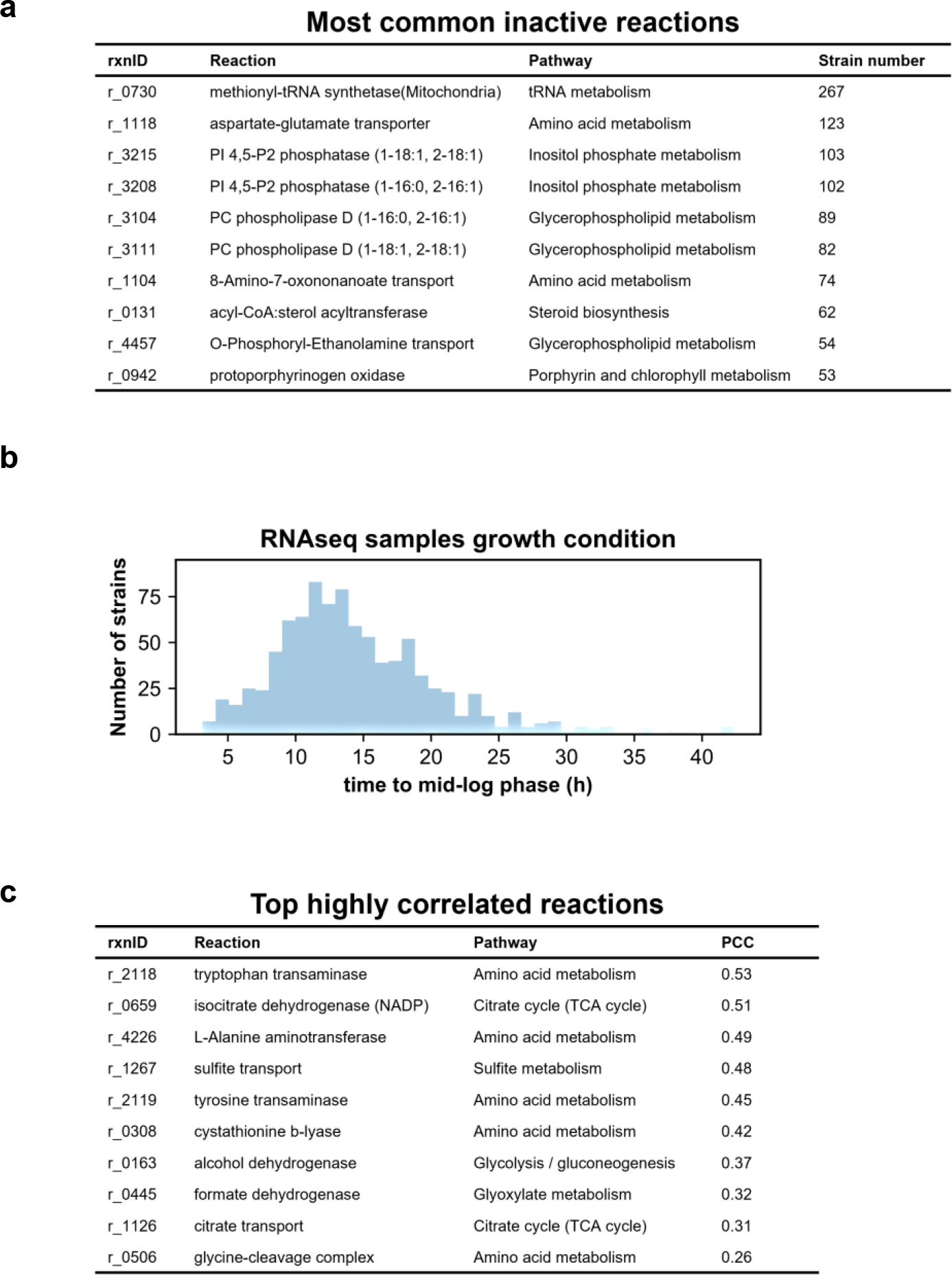
Characterization of important metabolic reactions associated with transcriptional regulation. Annotation of frequently inactive reactions attributed to relatively low expression levels of associated genes (a). Evaluation of relative growth rates for 969 RNA-seq samples (b). Annotations of the top 10 reactions ranked by correlation coefficients between absolute flux values and gene transcription levels (c).

**Supplementary Figure 5.**
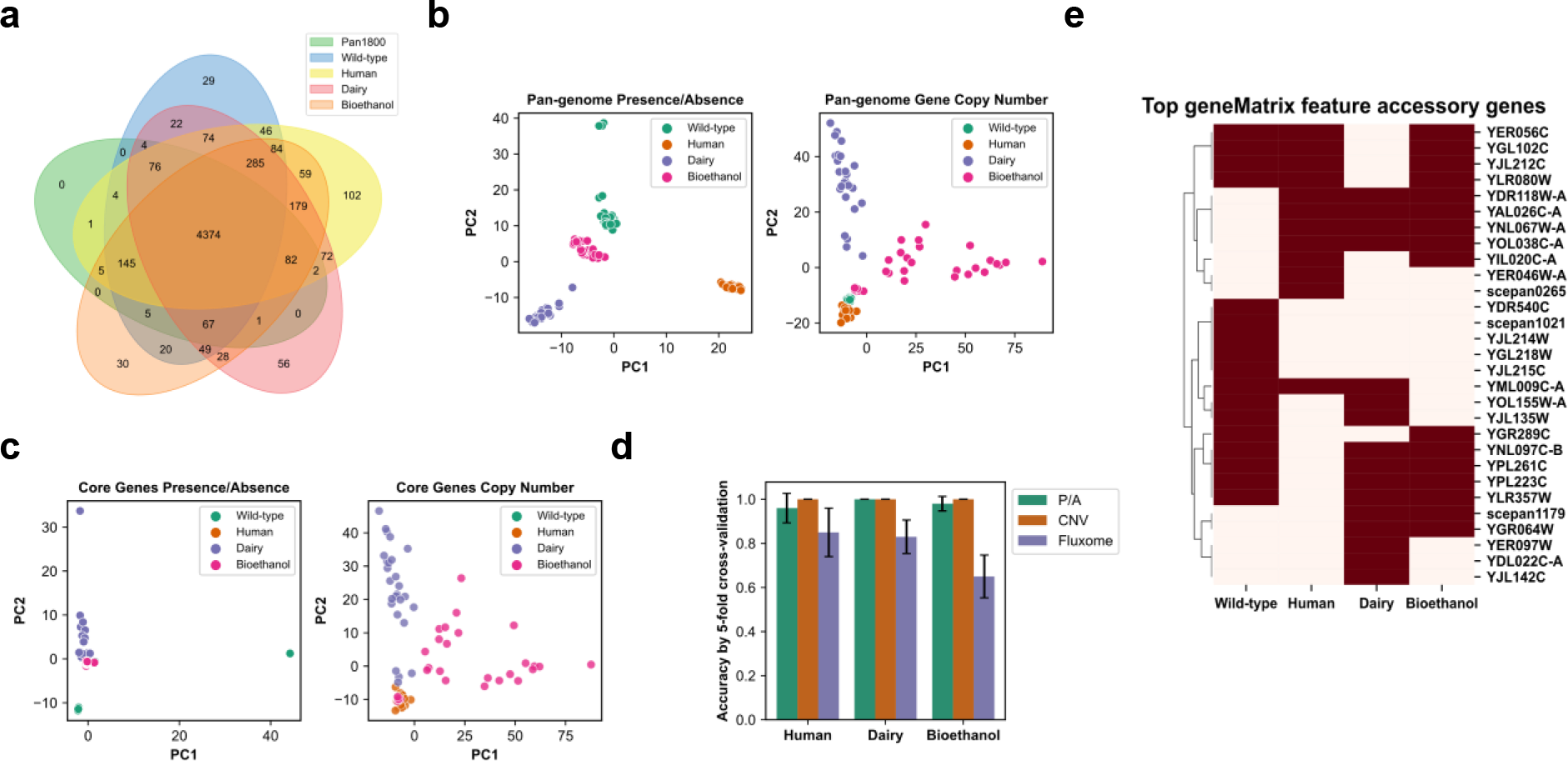
Pan-genome analysis for human, dairy, bioethanol, and wild-type clades. Core genome comparison among bioethanol, human, dairy, and wild-type strains (a). PCA clustering analysis based on the gene presence/absence matrix and the gene copy number matrix defined in the pan-genome for strains from the above 4 clades (b). PCA clustering analysis based on the gene presence/absence matrix and the gene copy number matrix defined in the core genome for strains from the above 4 clades (c). Accuracy scores obtained by 5-fold cross-validation with a random forest classifier using different parameters as inputs. P/A: gene presence/absence matrix; CNV: gene copy number content; Fluxome: metabolic flux profile (d). Top gene features from gene presence/absence matrix used to classify strains from the above 4 clades by a random forest algorithm (e).

**Supplementary Figure 6.**
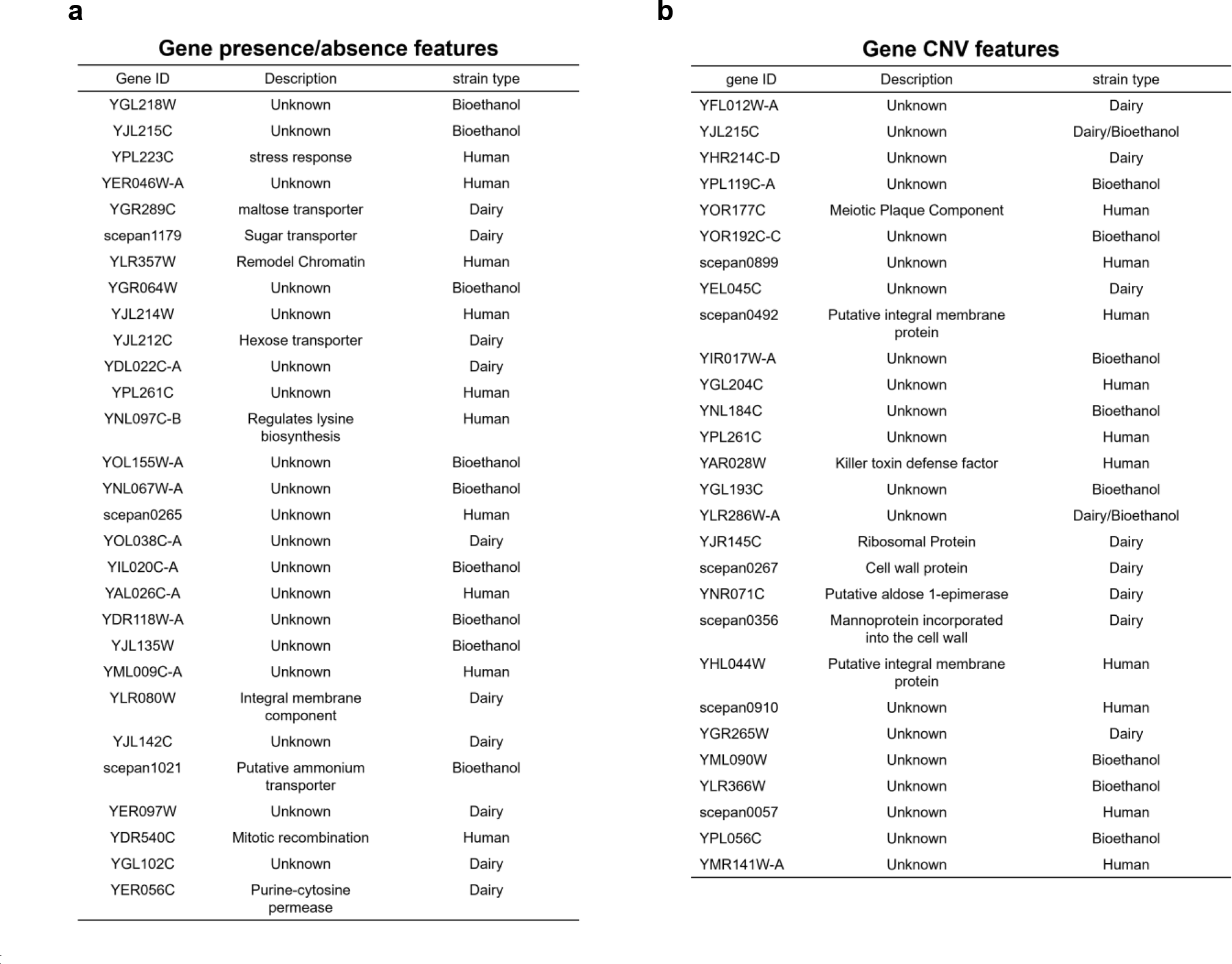
Functional annotations of genetic features for human, dairy, bioethanol, and wild-type clades. Functional annotations of top gene Presence/Absence features to classify human, dairy, bioethanol, and wild-type strains by a random forest algorithm (a). Functional annotations of top gene copy number variation features to classify human, dairy, bioethanol, and wild-type strains by random forest algorithm (b).

**Supplementary Figure 7.**
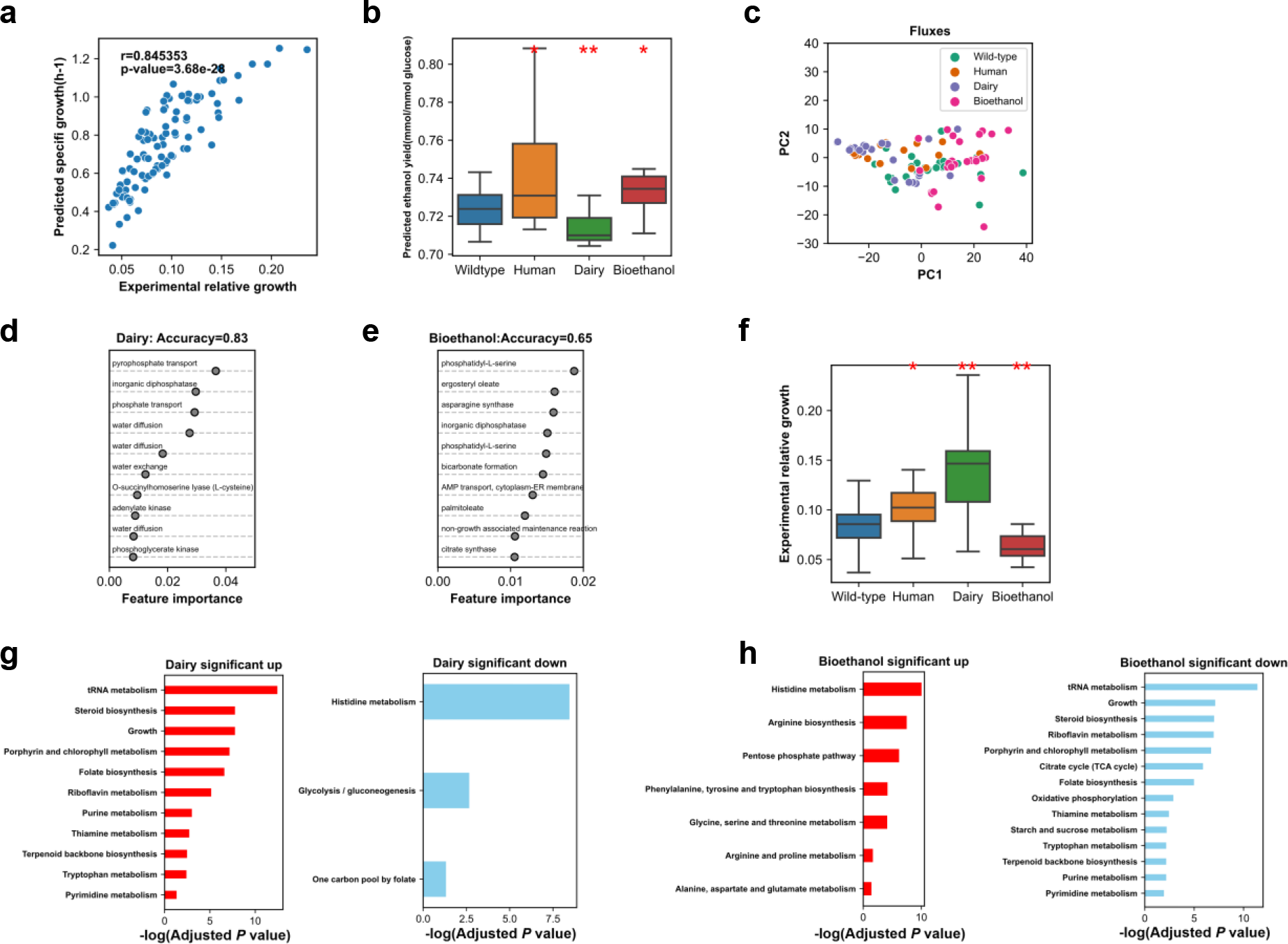
Comparative metabolic network analysis of human, dairy, and bioethanol clades. Correlation evaluation between the relative growth rata (Estimated based on measured t-mid time) and the corresponding simulated growth rate based on ssGEMs (a). Predicted ethanol production across human, dairy, bioethanol and wild-type clades (b). Statistical significance was determined using a two-sided T-test, significance denoted as *: *P* value < 0.05 and **: *P* value < 0.01. PCA clustering analysis of genome-scale flux distributions for human, dairy, bioethanol, and wild-type strains (c). Top metabolic flux features used to classify dairy strains and wild-type by a random forest algorithm (d). Top metabolic flux features used to classify bioethanol strains and wild-type by a random forest algorithm (e). Growth rate comparison across human, dairy, bioethanol, and wild-type clades based on experimental growth rate with glucose as the main carbon source (estimated based on the measured t-mid time dataset) (f). Enrichment analysis of reactions with differential metabolic fluxes between diary-related strains and the wild-type. (g). Enrichment analysis of reactions with differential metabolic fluxes between dairy-related strains and the wild-type (h).

**Supplementary Figure 8.**
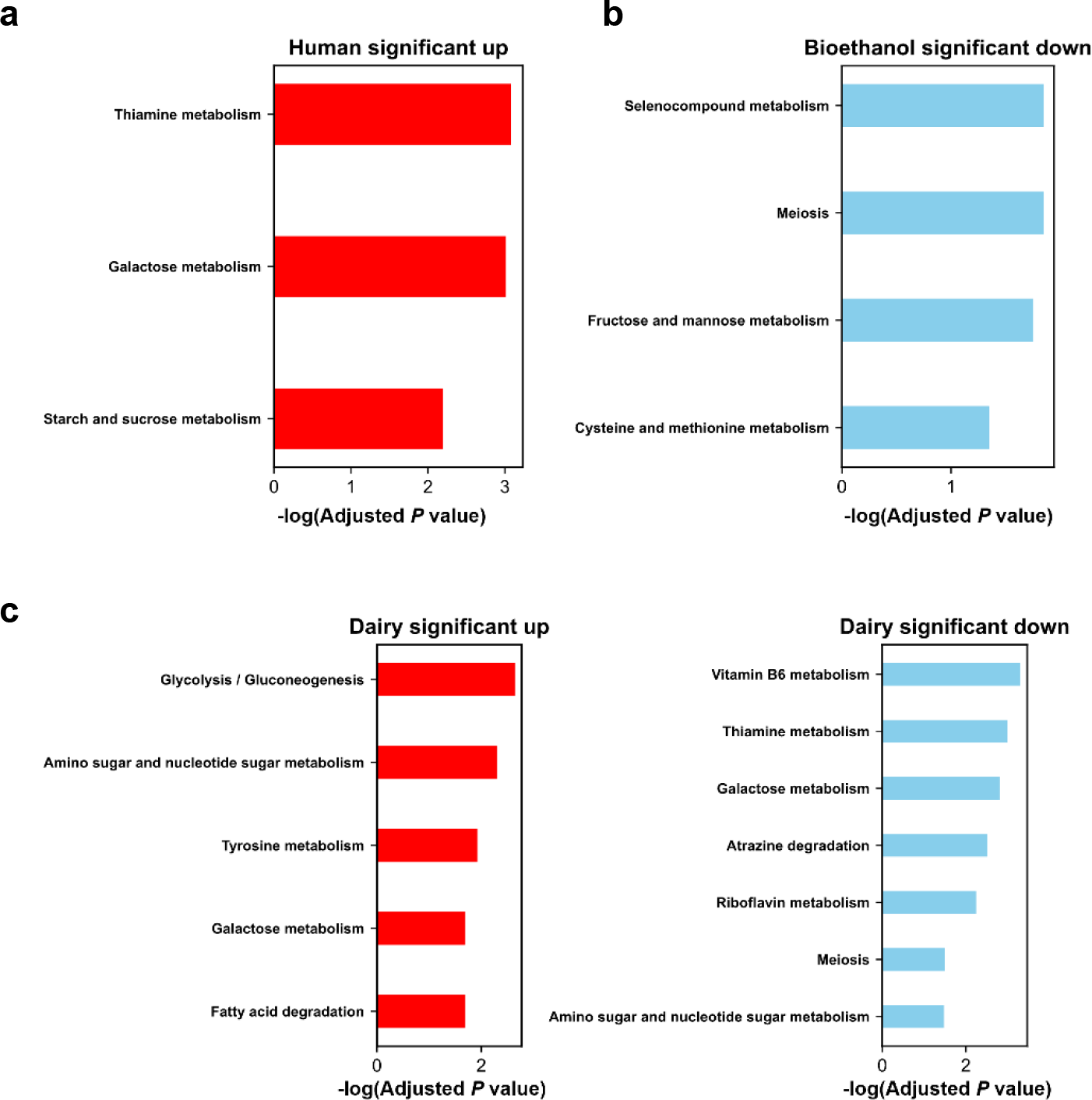
KEGG pathway enrichment analysis of differentially expressed genes in the human (a), bioethanol (b) and dairy (c) clades.

## Reference

1. Steensels, J., Gallone, B., Voordeckers, K. & Verstrepen, K. J. Domestication of Industrial Microbes. Current Biology 29, R381–R393 (2019).

2. Nielsen, J. Yeast cell factories on the horizon. Science 349, 1050–1051 (2015).

3. Goffeau, A. et al. Life with 6000 Genes. Science 274, 546–567 (1996).

4. Botstein, D. & Fink, G. R. Yeast: an experimental organism for 21st Century biology. Genetics 189, 695–704 (2011).

5. Shao, Y. et al. Creating a functional single-chromosome yeast. Nature 560, 331–335 (2018).

6. Winzeler, E. A. et al. Functional Characterization of the *S. cerevisiae* Genome by Gene Deletion and Parallel Analysis. Science 285, 901–906 (1999).

7. Giaever, G. et al. Functional profiling of the Saccharomyces cerevisiae genome. Nature 418, 387– 391 (2002).

8. Turco, G. et al. Global analysis of the yeast knockout phenome. Science Advances (2023).

9. Peter, J. et al. Genome evolution across 1,011 Saccharomyces cerevisiae isolates. Nature 556, 339– 344 (2018).

10. Marsit, S. et al. Evolutionary biology through the lens of budding yeast comparative genomics. Nat Rev Genet 18, 581–598 (2017).

11. Zhu, Y. O., Sherlock, G. & Petrov, D. A. Whole Genome Analysis of 132 Clinical Saccharomyces cerevisiae Strains Reveals Extensive Ploidy Variation. G3 (Bethesda) 6, 2421–34 (2016).

12. Lee, T. J. et al. Extensive sampling of *Saccharomyces cerevisiae* in Taiwan reveals ecology and evolution of predomesticated lineages. Genome Res. genome;gr.276286.121v2 (2022) doi:10.1101/gr.276286.121.

13. Gallone, B. et al. Domestication and Divergence of Saccharomyces cerevisiae Beer Yeasts. Cell 166, 1397–1410 e16 (2016).

14. Duan, S. F. et al. The origin and adaptive evolution of domesticated populations of yeast from Far East Asia. Nat Commun 9, 2690 (2018).

15. Díaz-Muñoz, C., Verce, M., De Vuyst, L. & Weckx, S. Phylogenomics of a Saccharomyces cerevisiae cocoa strain reveals adaptation to a West African fermented food population. iScience 25, 105309 (2022).

16. Kang, K. et al. Linking genetic, metabolic, and phenotypic diversity among Saccharomyces cerevisiae strains using multi-omics associations. Gigascience 8, (2019).

17. Han, D. Y. et al. Adaptive Gene Content and Allele Distribution Variations in the Wild and Domesticated Populations of Saccharomyces cerevisiae. Front Microbiol 12, 631250 (2021).

18. Caudal, É. et al. Pan-transcriptome reveals a large accessory genome contribution to gene expression variation in yeast. Nat Genet (2024) doi:10.1038/s41588-024-01769-9.

19. Vernikos, G., Medini, D., Riley, D. R. & Tettelin, H. Ten years of pan-genome analyses. Current Opinion in Microbiology 23, 148–54 (2015).

20. Li, G., Ji, B. & Nielsen, J. The pan-genome of *Saccharomyces cerevisiae*. FEMS Yeast Research 19, foz064 (2019).

21. O’Brien, E. J., Monk, J. M. & Palsson, B. O. Using Genome-scale Models to Predict Biological Capabilities. Cell 161, 971–987 (2015).

22. Förster, J., Famili, I., Fu, P., Palsson, B. Ø. & Nielsen, J. Genome-Scale Reconstruction of the Saccharomyces cerevisiae Metabolic Network. Genome Res 13, 244–253 (2003).

23. Chen, Y., Li, F. & Nielsen, J. Genome-scale modeling of yeast metabolism: retrospectives and perspectives. FEMS Yeast Res (2022) doi:10.1093/femsyr/foac003.

24. Simão, F. A., Waterhouse, R. M., Ioannidis, P., Kriventseva, E. V. & Zdobnov, E. M. BUSCO: assessing genome assembly and annotation completeness with single-copy orthologs. Bioinformatics 31, 3210–3212 (2015).

25. Cantalapiedra, C. P., Hernández-Plaza, A., Letunic, I., Bork, P. & Huerta-Cepas, J. eggNOG-mapper v2: Functional Annotation, Orthology Assignments, and Domain Prediction at the Metagenomic Scale. Molecular Biology and Evolution 38, 5825–5829 (2021).

26. Yuan, Y. et al. Pan-Genome Analysis of Transcriptional Regulation in Six Salmonella enterica Serovar Typhimurium Strains Reveals Their Different Regulatory Structures. mSystems e00467–22 (2022) doi:10.1128/msystems.00467-22.

27. Yang, T. & Gao, F. High-quality pan-genome of *Escherichia coli* generated by excluding confounding and highly similar strains reveals an association between unique gene clusters and genomic islands. Briefings in Bioinformatics bbac283 (2022) doi:10.1093/bib/bbac283.

28. Norsigian, C. J. et al. Systems biology approach to functionally assess the Clostridioides difficile pangenome reveals genetic diversity with discriminatory power. PNAS 119, e2119396119 (2022).

29. Shen, X. X. et al. Tempo and Mode of Genome Evolution in the Budding Yeast Subphylum. Cell 175, 1533–1545 e20 (2018).

30. Lu, H. et al. Yeast metabolic innovations emerged via expanded metabolic network and gene positive selection. Molecular Systems Biology 17, (2021).

31. Riley, R. et al. Comparative genomics of biotechnologically important yeasts. PNAS 113, 9882–7 (2016).

32. Lu, H. et al. A consensus S. cerevisiae metabolic model Yeast8 and its ecosystem for comprehensively probing cellular metabolism. Nat Commun 10, 3586 (2019).

33. Chen, S., Brockenbrough, J. S., Dove, J. E. & Aris, J. P. Homocitrate Synthase Is Located in the Nucleus in the Yeast Saccharomyces cerevisiae. J Biol Chem 272, 10839–10846 (1997).

34. Isogai, S. et al. High-Level Production of Lysine in the Yeast Saccharomyces cerevisiae by Rational Design of Homocitrate Synthase. Applied and Environmental Microbiology 87, e00600–21 (2021).

35. Opdam, S. et al. A Systematic Evaluation of Methods for Tailoring Genome-Scale Metabolic Models. Cell Systems 4, 318–329.e6 (2017).

36. Culley, C., Vijayakumar, S., Zampieri, G. & Angione, C. A mechanism-aware and multiomic machine-learning pipeline characterizes yeast cell growth. PNAS 117, 18869–18879 (2020).

37. Becker, S. A. & Palsson, B. O. Context-Specific Metabolic Networks Are Consistent with Experiments. PLOS Computational Biology 4, e1000082 (2008).

38. Li, Y., Holmes, W. B., Appling, D. R. & RajBhandary, U. L. Initiation of Protein Synthesis in Saccharomyces cerevisiae Mitochondria without Formylation of the Initiator tRNA. Journal of Bacteriology 182, 2886–2892 (2000).

39. Wiltrout, E., Goodenbour, J. M., Fréchin, M. & Pan, T. Misacylation of tRNA with methionine in Saccharomyces cerevisiae. Nucleic Acids Res 40, 10494–10506 (2012).

40. Jiang, Y.-Q. & Lin, J.-P. Recent progress in strategies for steroid production in yeasts. World J Microbiol Biotechnol 38, 93 (2022).

41. Henry, S. A., Kohlwein, S. D. & Carman, G. M. Metabolism and Regulation of Glycerolipids in the Yeast Saccharomyces cerevisiae. Genetics 190, 317–349 (2012).

42. Ye, C., Bandara, W. M. M. S. & Greenberg, M. L. Regulation of Inositol Metabolism Is Fine-tuned by Inositol Pyrophosphates in Saccharomyces cerevisiae♦. J Biol Chem 288, 24898–24908 (2013).

43. Gmelch, L. et al. Comprehensive Vitamer Profiling of Folate Mono- and Polyglutamates in Baker’s Yeast (Saccharomyces cerevisiae) as a Function of Different Sample Preparation Procedures. Metabolites 10, 301 (2020).

44. Hohmann, S. & Meacock, P. A. Thiamin metabolism and thiamin diphosphate-dependent enzymes in the yeast Saccharomyces cerevisiae: genetic regulation. Biochim Biophys Acta 1385, 201–219 (1998).

45. Thiaville, P. C., Iwata-Reuyl, D. & de Crécy-Lagard, V. Diversity of the biosynthesis pathway for threonylcarbamoyladenosine (t(6)A), a universal modification of tRNA. RNA Biol 11, 1529–1539 (2014).

46. Qiu, D. et al. Analysis of inositol phosphate metabolism by capillary electrophoresis electrospray ionization mass spectrometry. Nat Commun 11, 6035 (2020).

47. Dekker, W. J. C., Wiersma, S. J., Bouwknegt, J., Mooiman, C. & Pronk, J. T. Anaerobic growth of Saccharomyces cerevisiae CEN.PK113-7D does not depend on synthesis or supplementation of unsaturated fatty acids. FEMS Yeast Research 19, (2019).

48. Mercurio, K., Singh, D., Walden, E. & Baetz, K. Global analysis of Saccharomyces cerevisiae growth in mucin. G3 (Bethesda) 11, jkab294 (2021).

49. Li, G., Liu, L., Du, W. & Cao, H. Local flux coordination and global gene expression regulation in metabolic modeling. Nat Commun 14, 5700 (2023).

50. Chubukov, V. et al. Transcriptional regulation is insufficient to explain substrate-induced flux changes in Bacillus subtilis. Molecular Systems Biology 9, 709 (2013).

51. Bordel, S., Agren, R. & Nielsen, J. Sampling the Solution Space in Genome-Scale Metabolic Networks Reveals Transcriptional Regulation in Key Enzymes. PLOS Computational Biology 6, e1000859 (2010).

52. Yu, R., Vorontsov, E., Sihlbom, C. & Nielsen, J. Non-random organization of flux control mechanisms in yeast central metabolic pathways. 2021.12.15.472747 Preprint at 10.1101/2021.12.15.472747 (2021).

53. Hackett, S. R. et al. Systems-level analysis of mechanisms regulating yeast metabolic flux. Science 354, aaf2786–aaf2786 (2016).

54. Raghavan, V., Aquadro, C. F. & Alani, E. Baker’s Yeast Clinical Isolates Provide a Model for How Pathogenic Yeasts Adapt to Stress. Trends in Genetics 35, 804–817 (2019).

55. Lane, D. M., Valentine, D. L. & Peng, X. Genomic analysis of the marine yeast Rhodotorula sphaerocarpa ETNP2018 reveals adaptation to the open ocean. BMC Genomics 24, 695 (2023).

56. Engel, S. R. et al. New data and collaborations at the Saccharomyces Genome Database: updated reference genome, alleles, and the Alliance of Genome Resources. Genetics 220, iyab224 (2022).

57. Jenior, M. L., Jr, T. J. M., Dougherty, B. V. & Papin, J. A. Transcriptome-guided parsimonious flux analysis improves predictions with metabolic networks in complex environments. PLOS Computational Biology 16, e1007099 (2020).

58. Jacobus, A. P. et al. Comparative Genomics Supports That Brazilian Bioethanol Saccharomyces cerevisiae Comprise a Unified Group of Domesticated Strains Related to Cachaça Spirit Yeasts. Front. Microbiol. 12, (2021).

59. Niu, M. et al. ALR encoding dCMP deaminase is critical for DNA damage repair, cell cycle progression and plant development in rice. Journal of Experimental Botany 68, 5773–5786 (2017).

60. Yoshihisa, T., Ohshima, C., Yunoki-Esaki, K. & Endo, T. Cytoplasmic splicing of tRNA in Saccharomyces cerevisiae. Genes Cells 12, 285–297 (2007).

61. Jordá, T. & Puig, S. Regulation of Ergosterol Biosynthesis in Saccharomyces cerevisiae. Genes 11, 795 (2020).

62. Yang, P. et al. Thermotolerance improvement of engineered Saccharomyces cerevisiae ERG5 Delta ERG4 Delta ERG3 Delta, molecular mechanism, and its application in corn ethanol production. Biotechnol Biofuels Bioprod 16, 66 (2023).

63. Daignan-Fornier, B. & Pinson, B. Yeast to Study Human Purine Metabolism Diseases. Cells 8, 67 (2019).

64. Lacroute, F. Regulation of pyrimidine biosynthesis in Saccharomyces cerevisiae. J Bacteriol 95, 824–832 (1968).

65. Tengölics, R. et al. The metabolic domestication syndrome of budding yeast. Proceedings of the National Academy of Sciences 121, e2313354121 (2024).

66. Li, M. et al. Thiamine Biosynthesis in Saccharomyces cerevisiae Is Regulated by the NAD+- Dependent Histone Deacetylase Hst1. Mol Cell Biol 30, 3329–3341 (2010).

67. Wu, Y., Li, B., Miao, B., Xie, C. & Tang, Y.-Q. Saccharomyces cerevisiae employs complex regulation strategies to tolerate low pH stress during ethanol production. Microb Cell Fact 21, 247 (2022).

68. Bosi, E. et al. Comparative genome-scale modelling of Staphylococcus aureus strains identifies strain-specific metabolic capabilities linked to pathogenicity. PNAS 113, E3801–9 (2016).

69. Rosconi, F. et al. A bacterial pan-genome makes gene essentiality strain-dependent and evolvable. Nat Microbiol 7, 1580–1592 (2022).

70. Wang, M. et al. Annotation of 2,507 Saccharomyces cerevisiae genomes. Microbiology Spectrum 0,.

71. O’Donnell, S. et al. Telomere-to-telomere assemblies of 142 strains characterize the genome structural landscape in Saccharomyces cerevisiae. Nat Genet 55, 1390–1399 (2023).

72. Lu, H., Kerkhoven, E. J. & Nielsen, J. Multiscale models quantifying yeast physiology: towards a whole-cell model. Trends Biotechnol (2021) doi:10.1016/j.tibtech.2021.06.010.

73. Holt, C. & Yandell, M. MAKER2: an annotation pipeline and genome-database management tool for second-generation genome projects. BMC Bioinformatics 12, 491 (2011).

74. Pracana, R., Priyam, A., Levantis, I., Nichols, R. A. & Wurm, Y. The fire ant social chromosome supergene variant Sb shows low diversity but high divergence from SB. Molecular Ecology 26, 2864– 2879 (2017).

75. Cantalapiedra, C. P., Hernández-Plaza, A., Letunic, I., Bork, P. & Huerta-Cepas, J. eggNOG- mapper v2: Functional Annotation, Orthology Assignments, and Domain Prediction at the Metagenomic Scale. Molecular Biology and Evolution 38, 5825–5829 (2021).

76. Gopalakrishnan, S. et al. Guidelines for extracting biologically relevant context-specific metabolic models using gene expression data. Metabolic Engineering 75, 181–191 (2023).

77. Bjorkeroth, J. et al. Proteome reallocation from amino acid biosynthesis to ribosomes enables yeast to grow faster in rich media. PNAS 117, 21804–21812 (2020).

78. Ebrahim, A., Lerman, J. A., Palsson, B. O. & Hyduke, D. R. COBRApy: COnstraints-Based Reconstruction and Analysis for Python. BMC Systems Biology 7, 74 (2013).

79. Fang, Z., Liu, X. & Peltz, G. GSEApy: a comprehensive package for performing gene set enrichment analysis in Python. Bioinformatics 39, btac757 (2023).

80. Smith, A. B. et al. Enterococci enhance Clostridioides difficile pathogenesis. Nature 611, 780–786 (2022).

81. Vanoni, M. et al. A modular model integrating metabolism, growth, and cell cycle predicts that fermentation is required to modulate cell size in yeast populations. bioRxiv 2023.11.25.568635 (2023) doi:10.1101/2023.11.25.568635.

82. Muzellec, B., Teleńczuk, M., Cabeli, V. & Andreux, M. PyDESeq2: a python package for bulk RNA-seq differential expression analysis. Bioinformatics 39, btad547 (2023).

83. Zeng, Z. et al. OmicVerse: A single pipeline for exploring the entire transcriptome universe. 2023.06.06.543913 Preprint at 10.1101/2023.06.06.543913 (2023).

84. Kanehisa, M., Furumichi, M., Sato, Y., Kawashima, M. & Ishiguro-Watanabe, M. KEGG for taxonomy-based analysis of pathways and genomes. Nucleic Acids Res 51, D587–D592 (2023).

